# Superinfection plays an important role in the acquisition of complex *Plasmodium falciparum* infections among female *Anopheles* mosquitoes

**DOI:** 10.1101/2022.12.23.521802

**Authors:** Sophie Bérubé, Betsy Freedman, Diana Menya, Joseph Kipkoech, Lucy Abel, Zena Lapp, Steve M. Taylor, Wendy Prudhomme O’Meara, Andrew A. Obala, Amy Wesolowski

## Abstract

Studies of human malaria infections with multiple, genetically distinct parasites have illuminated mechanisms of malaria transmission. However, few studies have used the genetic diversity in mosquito infections to understand how transmission is sustained. We identified likely human sources of mosquito infections from a longitudinal cohort in Western Kenya based on genetic similarity between parasites and the timing of infections. We found that several human infections were required to reconstitute each mosquito infection and that multiple parasite clones were likely transmitted from infected humans to mosquitoes in each bite, suggesting that superinfection and co-transmission occur simultaneously and are important mechanisms of transmission. We further investigated this using an individual human and mosquito simulation model and found that co-transmission alone was unlikely to reproduce the high complexity of mosquito infections. We concluded that the superinfection of mosquitoes likely plays an important, but under studied, role in sustaining moderate to high malaria transmission.

## 1. Introduction

Following decades of progress in reducing malaria burden ^1–4^, global malaria cases increased from 2019 to 2020.^5^ Reversing this trend and continuing to reduce malaria transmission demands both scaling up current control efforts and expanding the set of available control measures. Many existing approaches, such as indoor residual spraying and bed net distribution, aim to lower transmission by reducing contact between humans and mosquitoes.^6,7^ However, targeting these types of control measures to reduce malaria transmission efficiently requires an understanding of which circumstances lead to successful *Plasmodium falciparum* transmission events between humans and mosquitoes, especially in natural settings.^8,9^

An important feature of malaria transmission dynamics is the acquisition of, often complex, *P. falciparum* infections where more than one genetically distinct parasite composes an infection.^10,11^ Similar to humans, mosquitoes can acquire these complex infections from humans by two possible mechanisms: 1) co-transmission, defined as the transmission of multiple, genetically distinct parasite clones from a single feed on an single infectious human, and 2) superinfection, defined as the transmission of multiple, genetically distinct parasite clones from a series of infectious feeds on multiple humans. These phenomena are not mutually exclusive; each feed in a series may involve the co-transmission of multiple clones and the relative contribution of superinfection and co-transmission to complex mosquito infections is not fully understood. In human infections, a study of complex infections in Malawi analyzing single-cell genome sequencing suggested that co-transmission is a more common mechanism than superinfection.^12^ However, the short lifespan of female *Anopheles* mosquitoes of only one to two weeks in natural settings, limits the number of blood meals each mosquito can take, suggesting that co-transmission may also be a more common mechanism for mosquitoes to acquire complex infections.^13^ However, studies that source the blood meals of mosquitoes in natural settings often consist of multiple different human sources, suggesting that superinfection also plays an important role.^14,15^

Measuring the relative contributions of these two mechanisms in *P. falciparum* transmission among mosquitoes is undermined by difficulties associated with observing natural human-to-mosquito transmission events. Experimentally, human landing catches or mosquito feeding assays (direct or membrane based) provide some of the most informative measures of human-to-mosquito transmission events and have identified human factors associated with infectivity as well as heterogeneity in mosquito biting patterns.^16,17,18,19^ However, both experimental approaches fall short of replicating natural transmission in key ways. Studies revealed lower rates of human-to-mosquito infections in membrane feeding assays than in direct skin feeding assays.^20,21^ While direct skin feeding assays and human landing catches provide a more realistic replica of natural transmission, results obtained from these experiments about mosquito biting behavior can be biased due to human variation in attractiveness to mosquitoes.^22,23^ Furthermore, these studies are often designed to elucidate factors associated with human infectiousness or attractiveness to mosquitoes and therefore may not be well suited to determine likely mosquito biting behaviors that lead to onward malaria transmission from humans to mosquitoes.^24,25^ To overcome difficulties in acquiring data about mosquito bionomics and its role in malaria transmission, modeling approaches have been employed, but these models rarely investigate the biting patterns that allow mosquitoes to acquire infections and have thus far not teased apart the roles of mosquito superinfection and co-transmission in sustaining onward malaria transmission.^26–29^

Genotyped *P. falciparum* parasites from infected humans and wild-fed mosquitoes from the same area and time frame provide a rich data source with which to identify likely human-to-mosquito transmission events. Previously, these studies have largely focused on the characteristics of infected humans that were probabilistically matched to infected mosquitoes by sharing *P. falciparum* clones.^12,15,30,31^ Prior work^31^ on these data found higher complexity of infection in mosquitoes (median of 6 distinct haplotypes per mosquito) as compared to humans (median of 3 distinct haplotypes per infection). The rich genetic information contained in high within-host diversity both on the mosquito and human level, allowed us to explore plausible mechanisms by which mosquitoes acquire infections. We used an algorithm to reconstitute the data on observed complex infections found in mosquito abdomens using parasites with shared genotypes found in human infections from a longitudinal cohort in a high transmission area of western Kenya. We then simulated patterns of mosquito feeding using an individual-based human and mosquito transmission model to identify likely mechanisms of transmission leading to the highly complex mosquito infections observed in these data. Using these data-informed models, we demonstrated that co-transmission alone is unlikely to produce mosquito infections as complex as those observed in this natural system suggesting that mosquito superinfection plays an important role in sustaining transmission in regions of moderate to high burden.

## 2. Methods

### 2.1 Sample Collection and molecular testing

A longitudinal cohort of 38 households in Bungoma County, Kenya, a region of high malaria transmission ^32,33^ was followed from June 2017 to July 2018. All household members one year of age and older were eligible for enrollment with a total of 268 individuals in the cohort. Dried blood spots (DBS), demographic and behavioral information were collected monthly. In addition to regular monthly, active followup, symptomatic visits were conducted with participants at the time of reported symptoms consistent with malaria infection, including diagnostic testing and DBS collection. A total of 902 asymptomatic infections (by PCR) were identified over 2312 monthly visits and a total of 137 symptomatic infections (also by PCR) were identified across 501 symptomatic visits (see Figure S1). Resting mosquitoes were collected from participating households by vacuum aspiration ^21^ weekly. The morphologically identified female *Anopheles* mosquitoes were transected to separate the abdomen from the head and thorax. Previously published sampling and laboratory procedures were used for these data^31,34^. Following genomic DNA extraction, both DBS and mosquito parts were tested for *P. falciparum* with a real-time PCR assay targeting the *P. falciparum* Pfr364 motif.

*P. falciparum* parasites identified by real-time PCR were sequenced across variable segments of the genes encoding the apical membrane antigen-1 (*Pfama1*) and the circumsporozoite protein (*Pfcsp*) using an Illumina MiSeq platform. Further details on PCR, read filtering, and haplotype calling, can be found in Sumner et al. (2019)^31^, and details on sequencing runs can be found in Figure S2. In total, across the 1039 human infections, 937 were successfully genotyped at the *Pfcsp* locus and 818 at the *Pfama1* locus. Among collected female *Anopheles* mosquito abdomens, 203 were positive for *P. falciparum* and a total of 185 infections found in mosquito abdomens were successfully genotyped at the *Pfcsp* locus and 177 at the *Pfama1* locus. In addition, 123 infections were identified in mosquito heads and were successfully genotyped *Pfcsp* and 121 at the *Pfama1* locus, although the focus of the analysis was on abdomen infections. After bioinformatic processing, a total of 209 unique *Pfama1* haplotypes were observed in human infections, 233 mosquito abdomens, and 79 in both populations, and 155 unique *Pfcsp* haplotypes were observed in human infections, 142 mosquito abdomens, and 44 in both populations.

### 2.2 Features of sequenced infections

For each infection, multiplicity of infection (MOI) was calculated independently for each sequenced locus as the total number of distinct *Pfcsp* or *Pfama1* haplotypes. Sumner et al. (2019)^31^ reported MOI distribution in humans that were lower than mosquito abdomens (see Figure S3); 32% of human infections were monoclonal (*Pfcsp* locus) compared to only 8% of mosquito abdomen infections. Results were broadly consistent for *Pfama1* (see Figure S4) and between mosquito heads and human infections (see Figure S5). Sensitivity analyses were performed on different subsets of haplotypes to, in part, account for potential technical effects across sequencing runs that could impact haplotype discovery and thus MOI (see Figure S2).

### 2.3 Reconstituting mosquito infections from human infections

To identify possible human-to-mosquito transmission events, parasite haplotypes observed in the abdomen of each mosquito (185 mosquitos) were reconstructed with haplotypes observed in human infections. The human infections considered eligible to contribute to the mosquito were those detected within the month before or after the mosquito was collected and had at least one *Pfcsp* or *Pfama1* haplotype in common with the mosquito’s infection. Of the eligible human infections, the ones considered likely matches for the mosquito were those that jointly reconstituted as many of the mosquito’s *Pfcsp* and *Pfama1* haplotypes as possible in the most parsimonious way, i.e. with the fewest possible number of human infections. It was not always possible to reconstitute all the haplotypes within a mosquito infection, and in this instance the most parsimonious option was still chosen.

A hypothetical matching scheme is outlined in Figure 1. Here a mosquito infection is comprised of haplotypes A, B and C (which could be a mix of *Pfcsp* and *Pfama1* haplotypes, since we initially matched on both loci jointly) with 3 eligible human infections (based on each having at least one of A, B and C haplotypes as well as being within one month of the mosquito sample point). These human infections can include overlapping and non-overlapping haplotypes with the mosquito infection (i.e. Human 1 is infected with haplotype A and E despite E not being found in the mosquito infection). Our matching scheme is based on finding the most parsimonious set of human infections to reconstitute the mosquito infections. In this example that would be both: Option M3 – humans 2 and 3 or option M4 – humans 1 and 3. These are equally parsimonious and fully reconstitute all the mosquito haplotypes (A, B, and C). In the instance of a tie like this one, we would randomly choose between these two options (M3 and M4). We would reject the other options because they would either involve more humans, as in option M2, or do not reconstitute the largest number of haplotypes within the mosquito infection, as in option M1 where the mosquito’s haplotype C is not accounted for in the matched human infections.

**Figure 1 :**
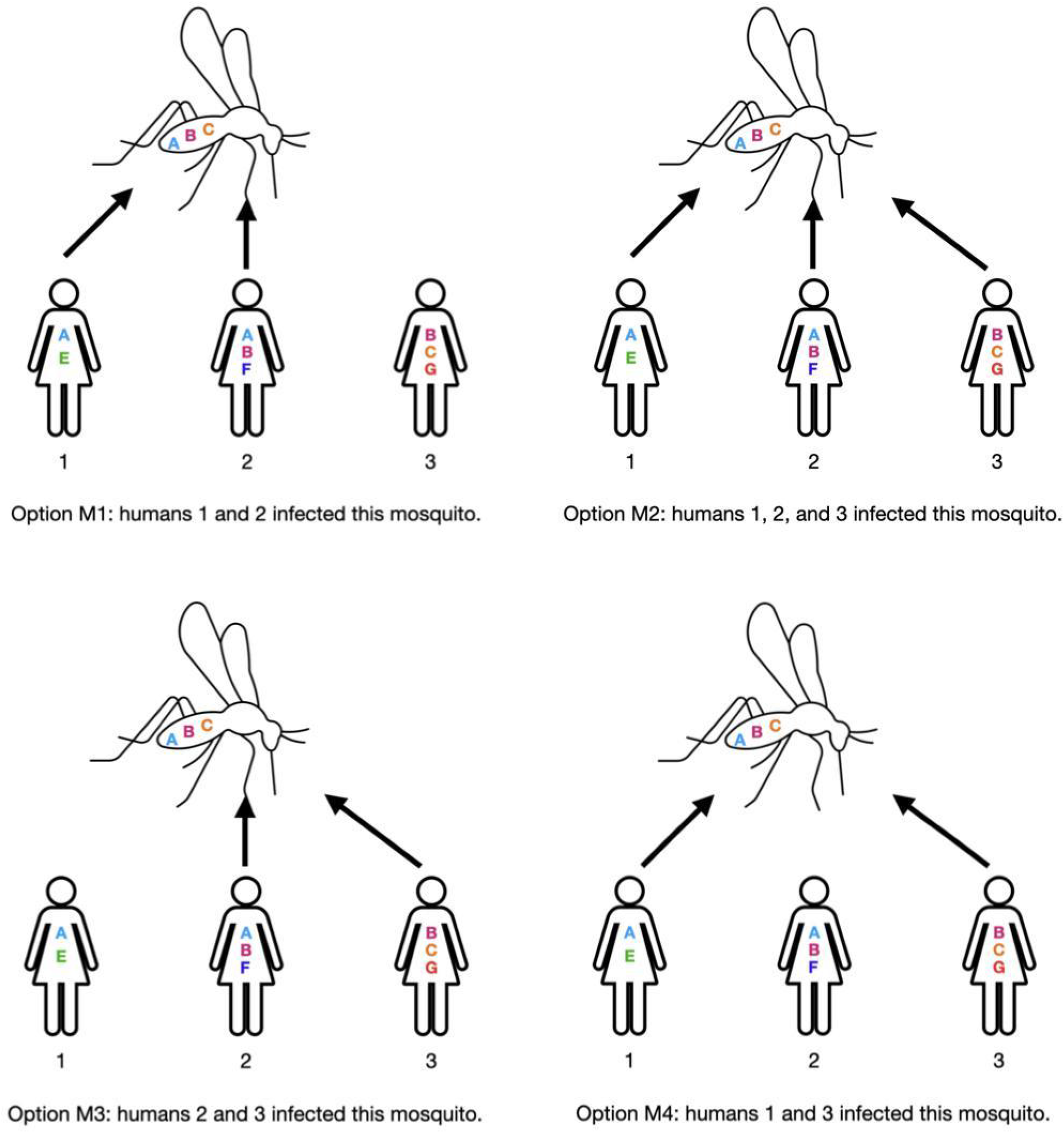
An example of possible human matches for a mosquito infection. Using our proposed method to reconstitute human-to-mosquito transmission events, we consider the most parsimonious set of human infections that will make up the mosquito infection. For a single mosquito infection (infected with haplotypes A, B, and C) the four possible options of human infections are shown as Options M 1-4. In this example, we would choose Option M3 or M4 since these strategies optimize the number of haplotypes reconstituted in the mosquito while minimizing the number of human infections, and hence feeds, needed.

This initial matching algorithm was described considering both loci jointly, however, we also reconstituted mosquito infections from human infections by considering each locus separately with similar results (see Figures S10-13). Furthermore, to account for potential systematic under or over discovery of haplotypes across samples of differing parasite density and sequencing read depth, we performed sensitivity analyses considering only common *Pfcsp* and *Pfama1* haplotypes jointly. Common haplotypes were defined as those with an amplicon specific population frequency in the top quartile of population frequencies (see Figure S8). We also considered unfiltered *Pfcsp* and *Pfama1* haplotypes jointly. Furthermore, to account for the fact that parsimonious matching may systematically underestimate rates of both mosquito and human superinfection we evaluated a matching algorithm where human matches were still within the same time windows but required sharing at least 2 or 3 haplotypes with the mosquito infection. We further implemented this matching algorithm based on 2 or 3 shared haplotypes separately for *Pfcsp* and *Pfama1* haplotypes.

To identify which factors are associated with human infections being in the set of parsimonious matches for a mosquito infection we used a multivariate logistic regression model. The outcome variable was whether or not a sampled human infection was in the set of parsimonious matches (see Section 2.3) for at least one mosquito infection. We included covariates at the person and infection level which were: whether the person reported sleeping under a net at the time of their sampled infection, the infection was symptomatic, the infection took place in the rainy or dry season, the person reported travel during the study period, the person had a symptomatic malaria infection during the study period, the total number of months the person was asymptomatically infected during the study period, the mean MOI of a person’s infections, and their age.

### 2.4 Malaria transmission simulation

A simulation model was developed to investigate realistic biological conditions that result in complex mosquito infections and was informed by the cohort data. An individual-based model of humans (n=200) and individual female *Anopheles* mosquitoes (stable population of 30,000) was constructed to simulate malaria transmission within our study population. To recapitulate the cohort data, we explicitly simulated infections at the individual haplotype level for a year (365 days) which is roughly the length of the cohort follow up period. We investigated the sensitivity of human and mosquito multiplicity of infection (MOI) and entomological inoculation rate (EIR) (measured as the number of infectious bites over 365 days) to various biological parameters that impact transmission, including mosquito biting patterns.

#### Initial conditions

We set initial conditions to broadly reflect those observed in the cohort. Initially, humans are randomly designated as infected with a probability approximately equal to the observed PCR positivity rate at monthly visits excluding sick visits (pr=0.3). If infected, the number of distinct haplotypes in the human infection (multiplicity of infection, MOI) is drawn from a truncated Poisson distribution fit to the data for the *Pfcsp* locus only (mean = 2, maximum = 16), data for the *Pfama1* was not used for calibration. Conditional on the initial MOI for a particular individual’s infection, the haplotypes that comprise that infection are drawn from the list of observed *Pfcsp* haplotypes with probability weights equal to the observed population frequency for *Pfcsp* haplotypes. Finally, the time since a person was initially infected with each haplotype present in their infection is drawn independently from a Poisson distribution with mean 20. This allows for a wide range of possible durations of human infections, and independent draws for each haplotype allow for the possibility of superinfection in humans.

The age-distribution of the initial population of 30,000 adult female *Anopheles* mosquitoes are drawn from a truncated Poisson distribution (mean = 4, maximum = 14 days) based on the daily survivorship of female *Anopheles* mosquitoes in the wild ^13^. Conditioned on age, the mosquito is designed as infected or not with older mosquitoes more likely to be infected and have more complex infections. Similar to human infections, the overall MOI distribution matches the distribution found in mosquito abdomens for *Pfcsp*. Like humans, the number of days since the mosquito was initially infected with each haplotype in their infection is drawn independently from a discrete uniform distribution between 1 and age of the mosquito.

#### Transmission dynamics

The model is incremented daily for a total of 722 days (357 days of burn-in and 365 days of sampling) where the following transitions can occur (Figure 2):

1. Mosquito biting behaviors: Whether a mosquito will take a human blood meal is determined probabilistically by the last time they fed. If they fed within the last three days, this is considered an “off day” and their probability of feeding on a human will be lower than on an “on day”, where the mosquito has not fed within the last three days. The exact probability of feeding on an “on” or “off” day is varied under different simulation scenarios (see Table 1). This setup allows, with relatively low probabilities, for mosquitoes to feed on humans multiple days in a row to complete a single blood meal. This is supported by evidence of female *Anopheles* mosquitoes imbibing multiple blood meals per gonotrophic cycle ^35,36^. This simulation also allows for the possibility that a mosquito may not feed on a human, which can occur if the mosquito feeds on other animals or does not successfully find a host. Though, in most cases the model assumes that a mosquito will feed on a human at least once in their lifetime, which is supported by the identification of 73% of the collected mosquitoes as *An. gambiae* or *An. funestus,* both highly anthropophilic species ^37,38^. Once a human feeding event has been determined, a discrete probability distribution is used to determine how many individuals the mosquito will feed on during the 24 hr period (see Table 2). All humans have an equal chance of being bitten.
2. Infectious bites: If a mosquito feeds on a human, and the human, mosquito, or both are infectious, a successful transmission event can occur. We consider different simulation scenarios (see Table 1) to investigate which haplotypes will be transmitted. Haplotypes are eligible to be transmitted from a human to a mosquito if they have been in the human for at least 14 days and vice versa if they have been in the mosquito for at least 9 days ^13^. Since we observed a wide range in the proportion of haplotypes shared (see Section 2.3, Figures 4B, S10B, S11B, S12B, and S13B) we allowed this to vary. Whether each eligible haplotype is transmitted during an infectious bite either from human to mosquito or vice versa is determined by independent draws from a Bernoulli distribution. We vary this probability under different simulation scenarios but within a scenario the value applies to all haplotypes.
3. Parasite clearance without treatment: In the cohort, we observe multiple instances of participants with asymptomatic infections who are subsequently PCR negative at a following time point without treatment (see Figure S14A). Additionally, we also observe instances when individual haplotypes were present at one asymptomatic visit, and absent at the next, while the person remained asymptomatically infected (see Figure S14B). Based on these observations, we allow for individual haplotypes to be cleared with some probability if the time since infection is 30 days or longer and vary this probability (see Table 1).
4. Symptomatic infections and treatment: Any infection containing at least one haplotype that was introduced in a human 7 or more days prior, can become symptomatic with a probability that was varied around values close to 0.1, the observed ratio of symptomatic to asymptomatic infections in the cohort (see Table1). We assume that symptomatic infections are sampled and treated immediately with all haplotypes in this infection cleared at the time of treatment, and that treatment offers protection from reinfection for 14 days ^39^.
5. Mosquito demography: for each mosquito, the probability of mosquito survival on any given day is negatively correlated with age. This relationship was chosen to produce a stable age-structured mosquito population with the maximum survival length to be 2 weeks (See Figure S9 for age distributions of mosquitoes). As mosquitoes die, they are replaced with newly emerged adult females to maintain a stable population size.

**Figure 2:**
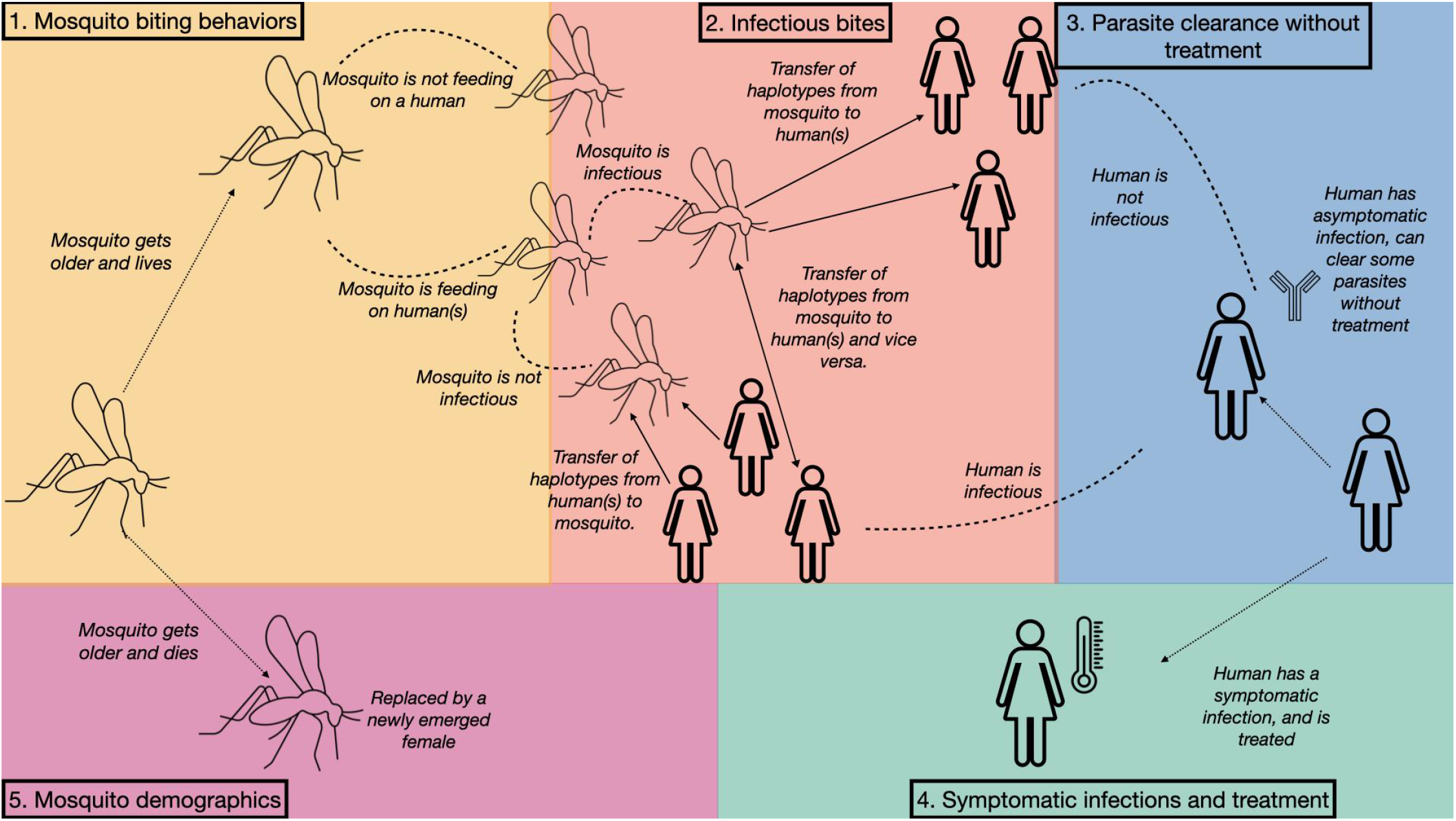
A graphic representation of the steps that take place during a single increment (one day) of the simulation. Complete descriptions of each stage (1-5) are detailed in Section 2.4.

**Figure 3:**
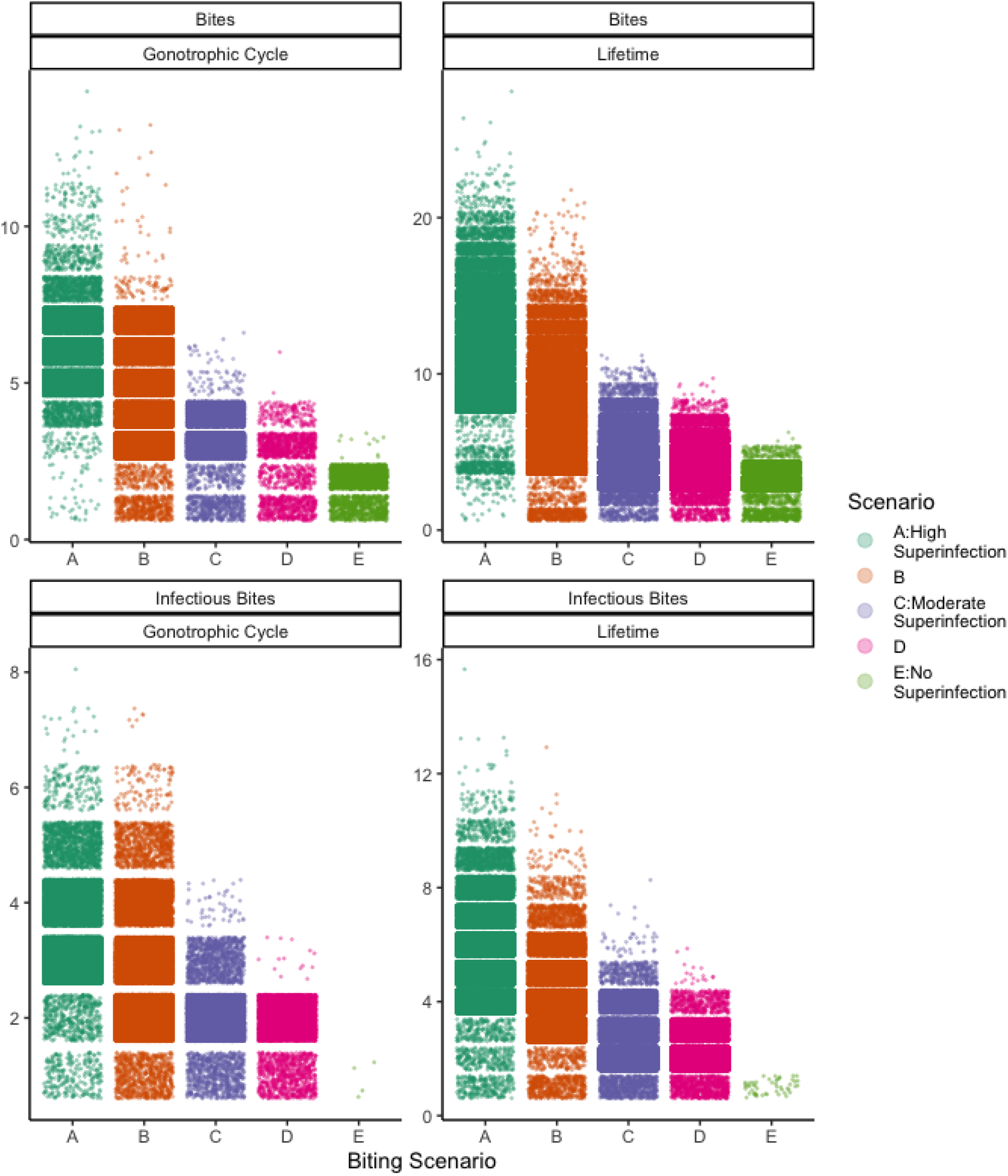
Number of bites and infectious bites taken by all 30,000 mosquitoes across 50 simulations in each of the five biting scenarios. Bites are counted both over the entire lifespan of a particular mosquito and over each gonotrophic cycle, which is assumed to last 3 days ^41–43^. An infectious bite is when at least one haplotype is transmitted between a human and a mosquito. In scenario E, superinfection (greater than 1 infectious bite in a single gonotrophic cycle or even over a mosquito’s lifetime) is not observed across 50 simulation scenarios each run for 365 days. However it is theoretically possible given that the total number of bites per gonotrophic cycle and lifetime for mosquitoes in scenario E can be greater than 1.

**Figure 4:**
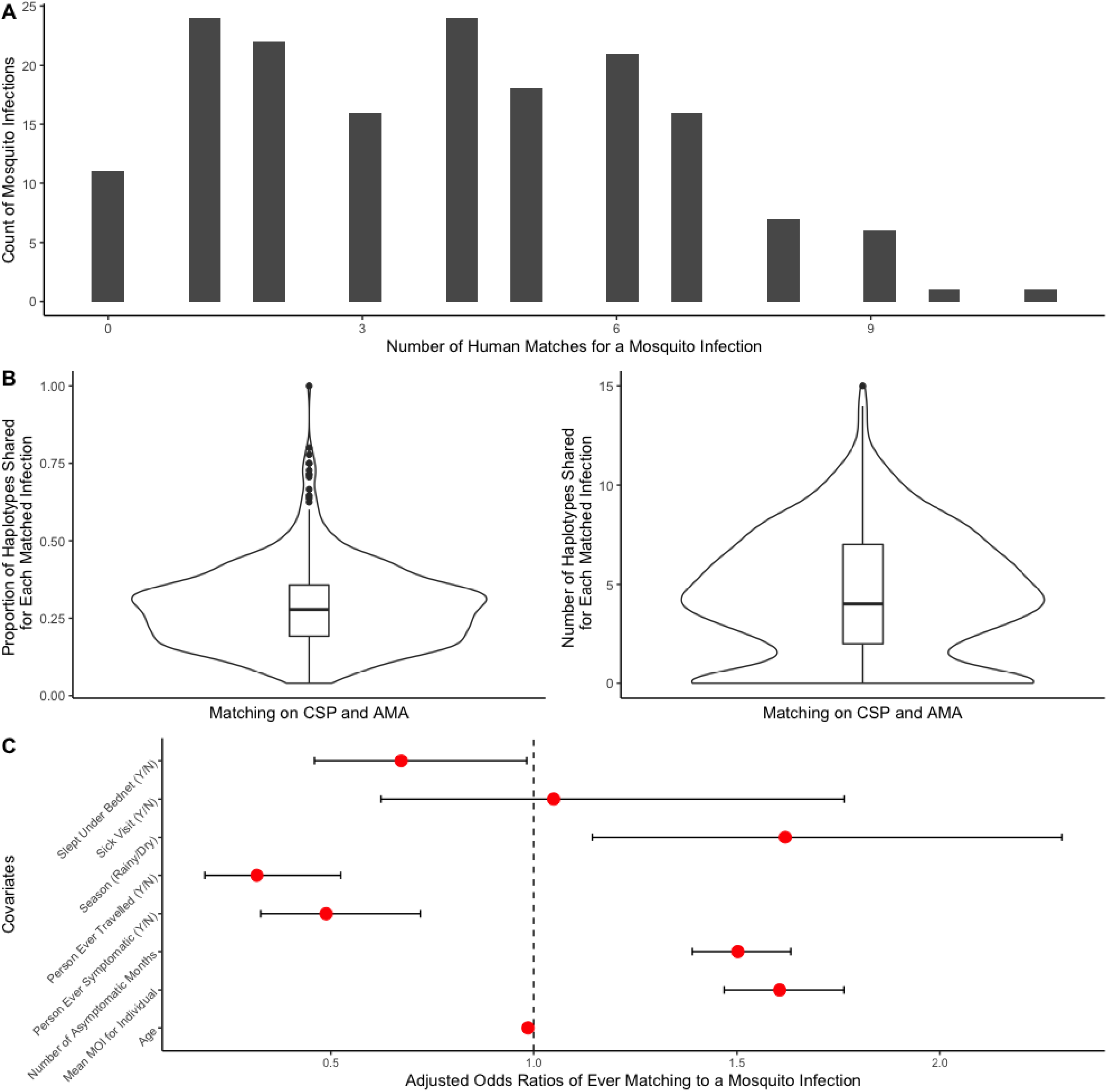
The Reconstituted mosquito infections from human infections considering both *Pfcsp* and *Pfama1*. A) The number of human infections needed to optimally reconstitute each mosquito infection considering both *Pfcsp* and *Pfama1* haplotypes simultaneously. B) The proportion and number of haplotypes from each human infection that are also in the matched mosquito abdomen infection. C) Results of a multiple logistic regression analyzing person and infection level factors that make a particular human infection more likely to reconstitute a mosquito infection. The outcome variable used was whether a particular sampled human infection was identified as reconstituting at least one mosquito infection.

**Table 1:** The values for parameters investigated for sensitivity analyses across simulations. Baseline values are those assigned to a parameter when it is not undergoing sensitivity analysis.

| Parameter | Baseline value | Range of values tested for sensitivity analyses |
| --- | --- | --- |
| Pr Symptomatic Infection | 0.05 | (0.01,0.3) |
| Pr Haplotype Clearing | 0.85 | (0.65,0.95) |
| Pr of Mosquito Feeding on On day | 0.1 | (0.01,0.3) |
| Pr of Mosquito Feeding on Off day | 0.01 | (0.001,0.05) |
| Pr of Transmission from Human to Mosquito | 0.6 | (0.2, 0.9) |
| Pr of Transmission from Mosquito to Human | 0.3 | (0.1,0.7) |

**Table 2:**
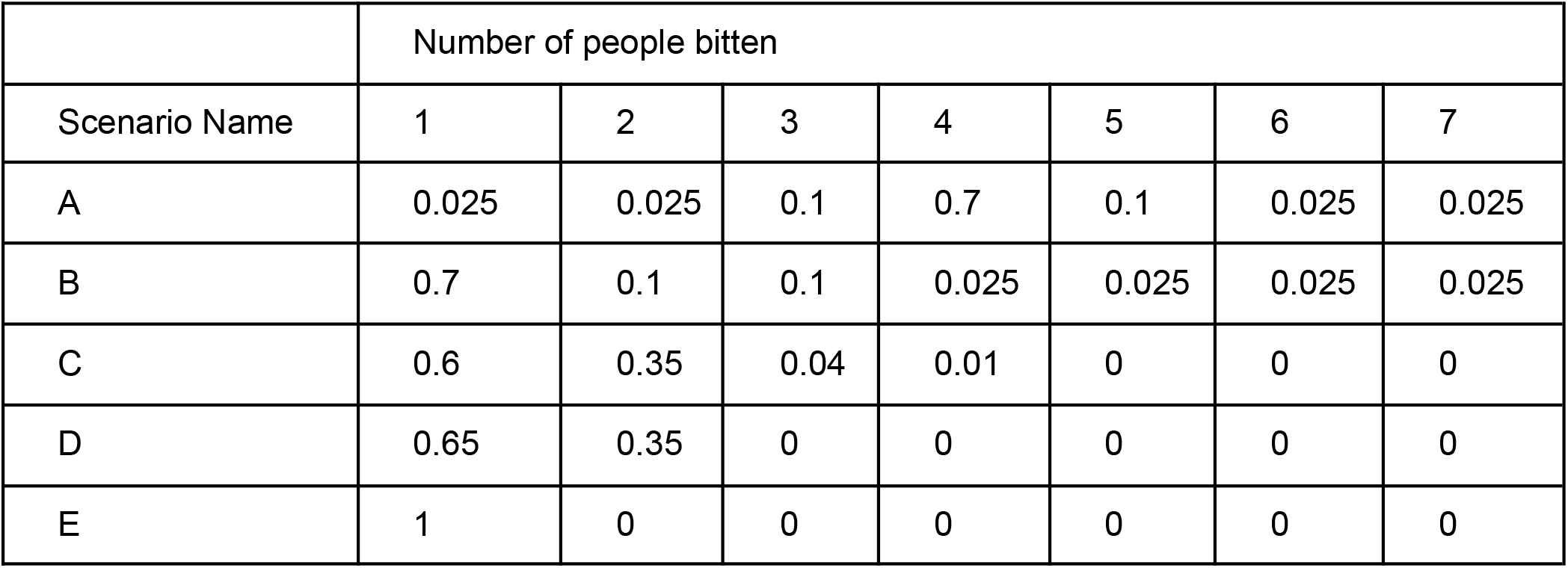
The mosquito biting patterns investigated in simulation. We considered a range of biting patterns where mosquitoes would feed on 1-7 individuals within a day. The five mosquito biting patterns (A-E) are shown with the probability of biting the number of individuals (columns). These patterns broadly reflect high (A/B), moderate (C/D) and low (E) biting scenarios. Scenarios A and B allow for a high probability of superinfection-within a 24 hour period-with more than two individuals, scenarios C and D allow for a moderate probability of superinfection-within a 24 hour period-with fewer individuals and scenario E allows for only co-transmission within a 24 hour period.

### 2.5 Sampling

Following a burn-in period (357 days), our simulation records human and mosquito infections sampled with the same frequency in the cohort. Mosquitos are sampled at random, weekly with population replacement (newly emerged adult females). All 200 humans are sampled monthly, and their symptomatic infections are all sampled similar to our sampling frame in the study.

### 2.6 Parameter values

To test the impact of certain parameters on the entomological inoculation rate and multiplicity of infection for human and mosquito infections, we performed a sensitivity analysis varying a single parameter at a time across 50 iterations. Table 1 shows the values tested for the investigated parameters (probability of symptomatic infection; probability of clearing a haplotype without treatment; “on” feeding day – probability of a mosquito biting a human if they have not fed in the last 3 days; “off” feeding day – the probability of a mosquito biting a human if they have fed in the last 3 days; probability of a haplotype being transmitted from a human to a mosquito and vice versa). The baseline parameter values during sensitivity analyses of other parameters are shown in Table 1 and were selected to be those values that produced the highest overlap with observed and published EIR and MOI values across all simulations (Figures 6, S17, S18, and S19). We also evaluated between-simulation stochasticity for these sensitivity analyses, which was generally low (see Figure S15).

**Figure 6:**
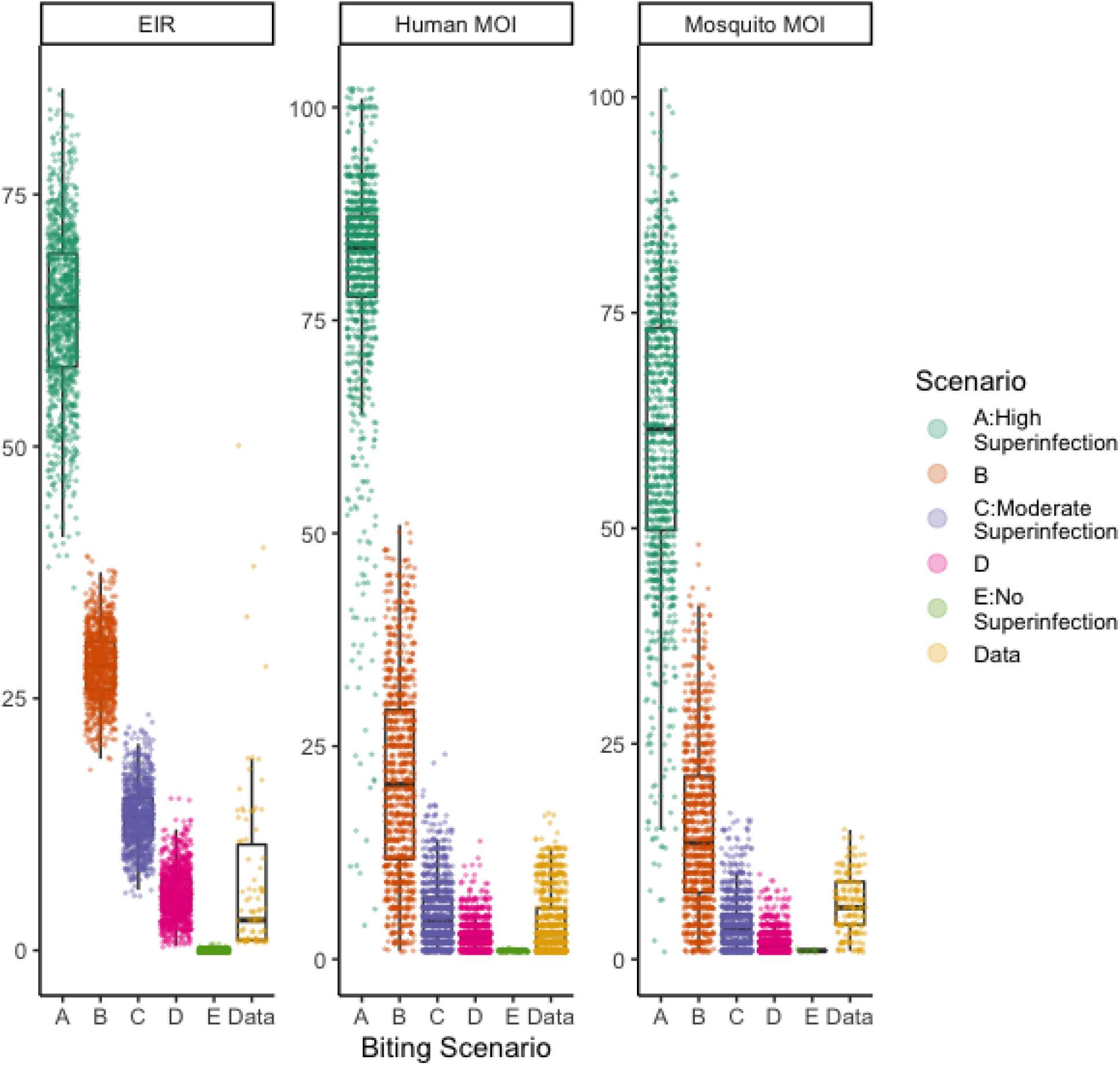
Entomological inoculation rate (EIR), and MOI under different mosquito biting patterns. Boxplots of EIR, human MOI, or mosquito MOI across simulations by biting scenario using the baseline values of all other simulation parameters (probability of haplotype clearance after 30 days; probability of a haplotype being transmitted from a mosquito to a human in an infectious bite; probability of a haplotype being transmitted from a human to a mosquito in an infectious bite; probability of feeding during an off day; probability of feeding during an on day; and probability of a symptomatic infection) for each simulation scenario. A random subset of 1000 points from the 50 simulation scenarios run for each biting pattern (A-E) are shown, all data points from the cohort are shown. EIR was calculated from the data by counting the number of infectious bites each person received during the study period. Here, a person is deemed to have an infectious bite from a mosquito if they are identified as a match using the parsimonious matching method described in Section 2.3. Published estimates of EIR for this region range from 16 to 24^45,46^.

We further investigated additional complexity in mosquito biting behavior. If a mosquito is feeding in a given 24-hour period, the number of people it will feed on during this time is determined based on the five scenarios shown below (see Table 2). We considered high biting scenarios (A) and low biting scenarios where the lowest biting scenario (E) only allows one human host per 24 hour feeding cycle. Higher biting scenarios (Scenarios A to B) make mosquito superinfection highly likely whereas the low biting scenarios (Scenarios D and E) make co-transmission more likely.

Finally, to account for potential heterogeneity in human attractiveness to mosquitoes and uneven distribution of infectious mosquito bites across people even within the same household ^24,40^, we performed some simulations where 20% of the population were *a priori* designated as more likely to be bitten.

Code for the simulation studies and matching algorithm can be found on the following GitHub repository: https://github.com/sberube3/mozzie_sim_manuscript/blob/main/.

## 3. Results

### 3.1 Multiple complex, asymptomatic human infections with high MOI values are required to reconstitute each mosquito infection

To investigate possible transmission dynamics leading to highly complex infections among mosquitoes, we performed a matching procedure where we reconstituted each infection observed in each mosquito abdomen with the minimum number of human infections required to explain the highest possible number of the mosquito’s haplotypes (see Section 2.3 for details). To reduce uncertainty in the time of infection for mosquitoes, we focused only on parasite genetic material (based on *Pfcsp* and *Pfama1* amplicons) found in abdomens since these are known to be more recently acquired than those found in the head^44^. Considering the amplicons jointly, results consistently demonstrated that more than one human infection was required to reconstitute a single mosquito infection with a median of 4 human infections required per mosquito infection (see Figure 4A, Figures S10A and S11A for results by amplicon). Only 11 (11/167, 5%) mosquito infections where both loci were successfully sequenced were unmatched to any human infection. Of these unmatched infections, 6 were monoclonal infections and the remaining 5 contained only rare haplotypes (population frequency in the bottom quartile of population frequencies). Matching results were consistent when only considering common haplotypes (see Figure S12A) or the unfiltered data (see Figure S13A). Furthermore, only 24 (14%) mosquito infections can be reconstituted with a single human infection (Figure 4A). Together these results suggest that superinfection – the process during which a mosquito feeds on multiple humans to acquire a complex infection – may be common in natural settings.

Using this matching scheme, we also explored the possible role of co-transmission, defined as the transmission of more than one genetically distinct parasite clone in a single infectious bite. We first estimated the proportion of haplotypes that were transmitted from a human infection successfully into its identified mosquito match during an infectious bite. The distribution of this proportion as well as the number of shared haplotypes across all identified human-mosquito matches considering both *Pfcsp* and *Pfama1* haplotypes is shown in Figure 4B. The median proportion of transmitted haplotypes from humans to mosquitoes is 0.35 and the mean number is 4, suggesting that, in many feeds upon people with polyclonal infections, mosquitoes may be acquiring more than one haplotype. Results were similar when considering each locus separately (see Figures S10B and S11B), for common haplotypes (see Figure S12B), and the unfiltered data (see Figure S13B).

We observed that, when considering both loci simultaneously, human infections were statistically significantly more likely to be in the set of parsimonious matches to a mosquito infection when they occurred during the rainy season (adjusted OR: 1.7; 95% CI: 1.2-2.4), in individuals with more asymptomatic infections (adjusted OR: 1.5; 95% CI: 1.4-1.6), and in individuals with higher MOI values (adjusted OR: 1.7; 95% CI: 1.5-1.8). Conversely, human infections were statistically significantly less likely to match to a mosquito infection for individuals who consistently slept under a bed net (adjusted OR: 0.6; 95% CI: 0.4-0.9), those who reported travel at any point during the study (adjusted OR: 0.3; 95% CI: 0.2-0.5), and from individuals who experienced at least one symptomatic infection (adjusted OR: 0.5; 95% CI: 0.4-0.8) (see Figure 4C). These results are generally unchanged when considering each locus separately, (Figures S10C and S11C), and across sensitivity analyses using common (Figure S12C) and unfiltered (Figure S13C) haplotypes. Symptom status, and MOI remain important predictors when humans that share 2 or 3 haplotypes with a mosquito’s infection are considered eligible for matching (Figure S16).

### 3.2 Mosquitoes are likely to feed on more than one individual within a 24 hour feeding period

The directly observed data suggest that both superinfection and co-transmission are important mechanisms for sustaining malaria transmission from humans to mosquitoes. To further investigate the roles of superinfection and co-transmission in mosquito infection, we carried out a series of individual based human and mosquito simulations using a simplified transmission model with parameters informed wherever possible by the cohort data (see Section 2.4). We compared the impact of different mosquito biting patterns on EIR and MOI distributions to evaluate whether superinfection, co-transmission, or a combination of both mechanisms were most likely to reproduce level of infection complexity and transmission intensity in the cohort data. We tested five different mosquito biting scenarios that each altered the number of humans a mosquito was likely to feed on in a single 24-hour period. Scenario A, which simulates a high rate of superinfection (i.e. a mosquito taking bloodmeals from more than one infected person in a 24 hour period) where the mosquito is likely to bite 4 or more people in a single 24-hour period is compared to scenarios with a decreasing likelihood of superinfection (scenarios B, C, and D). Ultimately, scenario E, which only allows for co-transmission within a 24 hour period and makes more than one infectious bite in a mosquitoes’ lifetime highly unlikely (see Figure 3) is considered the lowest biting scenario. Across multiple simulation parameter values (see Table 1), biting scenario A (high likelihood of superinfection) produces MOI values that are well above those observed in the data and EIR estimates well above those published for the region ^45,46^making four or more partial feeds during a 24 hour period an unlikely biting pattern for mosquitoes (see Figures 5 and S17). At the other extreme, across multiple simulation parameters, biting scenario E (making superinfection over a gonotrophic cycle or even a mosquito’s lifetime highly unlikely) produces MOI and EIR estimates that are well below those published and observed in the data and frequently leads to malaria extinction events, meaning that mosquito co-transmission alone is unlikely to sustain the levels of malaria transmission observed in the cohort (see Figure 5 and S17). Biting scenario C, which allows for a mosquito to bite up to 4 humans in a single 24-hour period and makes it likely the mosquito will bite between one and three infectious people per gonotrophic cycle (see Figure 3), produced the EIR and MOI distributions that most frequently overlapped with the observed and published estimates across multiple simulation parameters, suggesting that this rate of superinfection could sustain the levels of malaria transmission observed in the cohort (see Figure 5 and S17). Other biting scenarios that allowed for moderate amounts of superinfection (scenarios B and D) also produced MOI and EIR values in line with those observed in the data and literature, however, scenario C had the greatest overlap across all tested simulation parameters making it the most likely biting scenario (see Figures S18 and S19).

**Figure 5:**
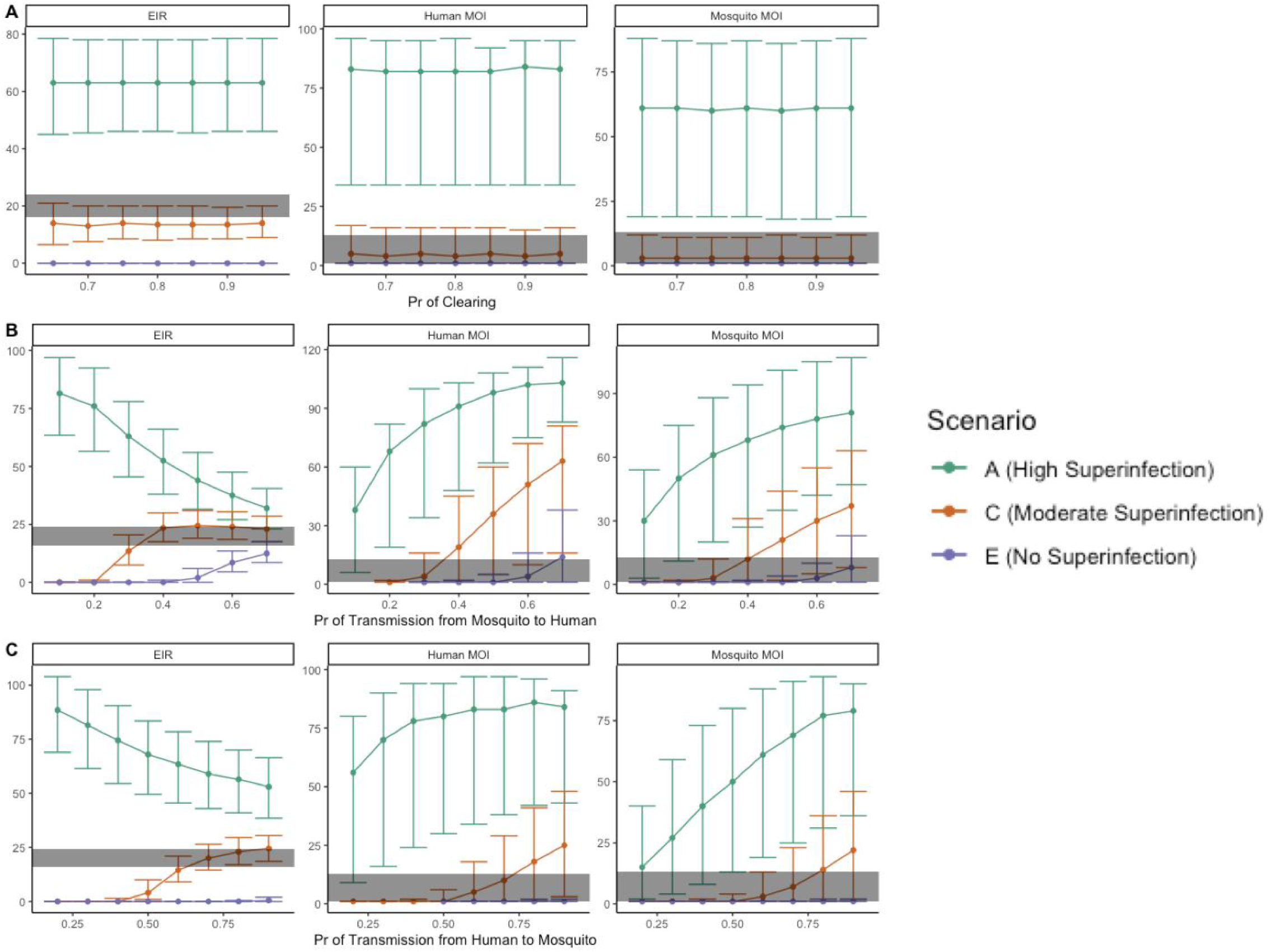
The sensitivity of entomological inoculation rate (EIR), human and mosquito multiplicity of infection (MOI) under different simulation parameters and biting scenarios. For multiple simulations, we varied parameters including (A) probability of haplotype clearance after 30 days; (B) probability of a haplotype being transmitted from a mosquito to a human in an infectious bite; and (C) probability of a haplotype being transmitted from a human to a mosquito in an infectious bite to investigate the impact of EIR, human MOI, and mosquito MOI values (median and 2.5th - 97.5th percentiles shown). Using three biting scenarios: high probability of superinfection (Scenario A), medium probability of superinfection (Scenario C), and no probability of superinfection (Scenario E), we find that high biting and low biting scenarios produce unrealistic estimates compared to the observed data (shown in the gray band represents the 2.5th and 97.5th percentile of the human and mosquito MOIs observed in the data and the range of published EIR estimates for the cohort area (16-24 bites per person, per year).

To compare EIR and MOI more directly under the different biting scenarios, we fixed values of the six simulation parameters at values that produced the highest overlap with observed and published EIR and MOI values across all five biting scenarios (A-E) (see Table 2). Using these parameters, scenario C produced the MOIs and EIR that most closely resembled published estimates and those observed in the data (see Figure 6). In simulations where 20% of the population was designated as more likely to be bitten by mosquitoes, biting scenario D appeared to better match the EIR and MOI values in the data than scenario C, however, scenario E, which makes superinfection, even over a mosquito’s lifespan, highly unlikely was still deemed implausible (see Figure S20). Collectively, our model simulations are consistent with an important role for a moderate amount of superinfection and co-transmission in producing complex mosquito infections and sustaining the levels of transmission observed in the cohort.

## 4. Discussion

A deeper understanding of malaria transmission can ultimately help us improve our approach to developing more effective control measures. Parasite genomics provide additional data sources with which to identify likely transmission events since parasites with shared genotypes may represent proximate events in the transmission chain. Ultimately, the complete picture of transmission requires identifying infections in both humans and mosquitoes. Here, we explore infections from both populations using *P. falciparum* parasite genotype data collected as part of a longitudinal cohort in a high transmission region of western Kenya. Previously, when investigating human and mosquito infections with cognate parasite genotypes, Sumner et al. (2019) ^31^ had identified that the mosquito infections had a much higher multiplicity of infection than human infections. These infections could be the result of individual mosquitoes acquiring multiple parasites from a single blood meal (co-transmission), from multiple blood meals taken from more than one human (superinfection), or a combination of both. Observing exactly how these factors contribute to sustaining transmission is extremely difficult in a natural setting, since it requires observing wild mosquito biting behaviors. However, using a combination of parasite genotype data and a simple simulation model of infection dynamics, we explored the degree to which superinfection and co-transmission contribute to mosquitoes acquiring complex infections. As opposed to many approaches that rely on explaining human infections with those observed in mosquitoes, we used a matching algorithm to reconstitute infections observed in mosquito abdomens with those sampled from humans based on time of sampling and shared *Pfcsp* and *Pfama1* haplotypes. Our method of reconstitution revealed that on average, multiple human sources, each with complex infections, were required to reconstitute the genetic diversity found in mosquito infections. Further, these matches rarely included only a single haplotype with a median of 4 distinct haplotypes being shared. These findings suggest that mosquitoes acquire complex infections through both superinfection and co-transmission.

To further investigate these empirical findings, we used a simple, individual-based human and individual-mosquito model to explore the likelihood of various mosquito biting patterns. We compared our results to published EIR values from similar settings^45,46^and to the observed distributions of human and mosquito MOIs in the cohort data. We explored scenarios with varied likelihoods of biting multiple infectious humans per gonotrophic cycle. Scenarios where mosquitoes were likely to feed on between 1 and 3 infectious humans per gonotrophic cycle were most plausible with higher feeding rates producing human and mosquito MOI and EIR values that far exceed those observed in the cohort data and lower rates resulting in MOI and EIR values well below those observed in the data. Tanken together, the results of both our matching algorithm and our transmission model suggest that superinfection plays an important role in acquiring complex mosquito infections observed in a high transmission area of western Kenya.

While this only represents one way to identify likely human-to-mosquito transmission events, our results were broadly consistent with previous analyses of the same data focused on reconstituting human infections^31^. While the cohort had very few missed visits and overall high follow-up rates, it may be true that sampled mosquitoes acquired infections from infected individuals outside of the cohort, or that despite intensive monthly sampling of human infections some infections within the cohort were missed. Furthermore, despite *Pfcsp* and *Pfama1* being highly diverse loci, we are unable to phase these segments to construct a more complete picture of individual parasites infecting humans and mosquitoes. Additional genotyping approaches that allow for the calculation of more direct measures of parasite relatedness, such as the probability of identity by descent, to be calculated could help better understand which parasites are related by transmission, however many of these approaches are not well suited to the highly polyclonal infections observed in this cohort. Furthermore, despite intensive sampling, in an area with such a high force of infection, it is impossible to pinpoint the exact timing of observed infections in both humans and mosquitoes, therefore we cannot differentiate co-transmission and superinfection events with certainty.

We constructed a simulation model to be able to explicitly model individual human and individual mosquito infections within a population similar to the setting in western Kenya. This allowed us to follow individual infection events between mosquitoes and humans to better understand how complex infections could arise in the mosquito population. Our simulation model was informed by the data whenever possible but certain assumptions are oversimplifications of reality. For example, we did not allow for any household clustering in the simulation despite evidence in other settings of within-household clustering of genetically related parasites ^47^. We observed little spatial clustering in our data, our regression analysis when including a random household intercept suggested little variation at the household level (see Figure S21). Moreover, we simplified parasite clearance in the absence of treatment and used a discrete probability, although more sophisticated models exist ^48^. We also did not include seasonal transmission patterns or account for other potential drivers of transmission such as human mobility. In addition, the simulation was sensitive to the number of mosquitoes simulated in the population (Figure S22). While it is difficult to quantify the true mosquito population relative to the human population in the area since our mosquito sampling only represents a biased subset of the full mosquito population, further exploration of how the number of mosquitoes interacts with other varying parameters in the model would be beneficial.

Our understanding of malaria transmission is limited by our ability to monitor and investigate infection dynamics within mosquitoes and their behavior in an entirely natural setting. While parasite genotyping data and intensive sampling strategies can provide insight into mechanisms of infection patterns, these will still be limited by our incomplete sampling of the full set of transmission events in a population. Simulation models, when informed by detailed cohort data, have the ability to supplement our understanding of mosquito bionomics by investigating the biological plausibility of different mosquito behaviors. Given the high MOI distribution observed among mosquitoes collected in the cohort study, and considering mosquitoes’ short lifespan, our investigation of how mosquitoes acquire such genetically diverse *P. falciparum* infections could provide insights into human reservoirs of transmission and parasite genetic diversity. While the former has clear implications for targeting more efficient interventions, the latter also has important implications for malaria surveillance; a pillar of most control strategies.

In particular, malaria molecular surveillance studies often rely on measures of parasite genetic diversity and relatedness to identify differences in transmission intensity or to uncover transmission dynamics such as spatial foci of transmission, but these studies also focus heavily on genotyping parasites in human infections ^49–51^. This and other studies have demonstrated that mosquitoes are an important reservoir of parasite genetic diversity ^52^. Beyond characterizing the level of parasite diversity, this study further disentangles the roles of mosquito superinfection, acquiring complex infections through feeding on multiple infectious humans, and co-transmission, acquiring complex infections by acquiring more than one, genetically distinct, parasite clone in a single bite. In this study, we have shown that heterogeneous and complex biting patterns that involve substantial levels of both co-transmission and superinfection may play an important role in sustaining high levels of malaria transmission. This conclusion points to a need for integrating analyses of mosquito infections into molecular surveillance studies to better account for the important role of mosquito transmission dynamics in sustaining malaria transmission.

## Author Contributions

SB and AW conceived and performed the quantitative analysis, and wrote the paper. SB, WPO, SMT, and AW interpreted the findings. WPO, SMT, BF, AAO, JKK, JM and LA designed and carried out the cohort study. WPO, SMT, BF and ZL performed the genetic analysis. All authors reviewed and revised the manuscript.

## Funding and Acknowledgements

AW is supported by a Career Award at the Scientific Interface from the Burroughs Wellcome Fund, and by the National Institute of Allergy and Infectious Diseases (R01AI146849). SB, RA, and AW are all supported by the National Institute of Health Director’s New Innovator Award, grant number DP2LM013102-0 and by the National Institute of Allergy and Infectious Diseases (1R01A1160780-01). This work was supported by NIAID (R21AI126024 to WPO and R01AI146849 to WPO and SMT). We thank the field technicians in Webuye for their engagement with the study participants: I. Khaoya, L. Marango, E. Mukeli, E. Nalianya, J. Namae, L. Nukewa, E. Wamalwa, and A. Wekesa.

## Data Availability and Ethical Approvals

The sequence data analyzed during the current study are available from NCBI (PRJNA646940). The study was approved by the ethical review boards of Moi University (2017/36), Duke University (Pro00082000), and the University of North Carolina at Chapel Hill (19–1273).

