## Supplemental Figures for "Superinfection plays an important role in the acquisition of complex *Plasmodium falciparum* infections among female *Anopheles* mosquitoes"

December 23, 2022

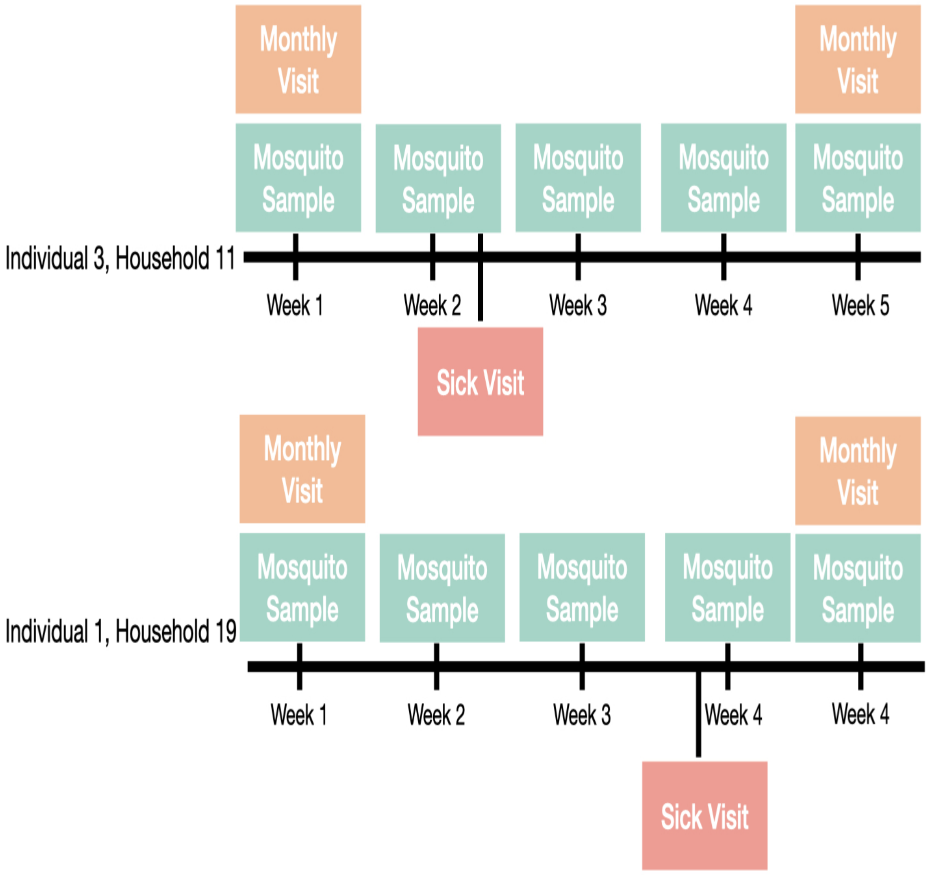


**Fig S 1:** Schematic depicting the sampling scheme for the cohort study in for two individuals in their respective households. Note that mosquito sampling occurs in each household weekly.

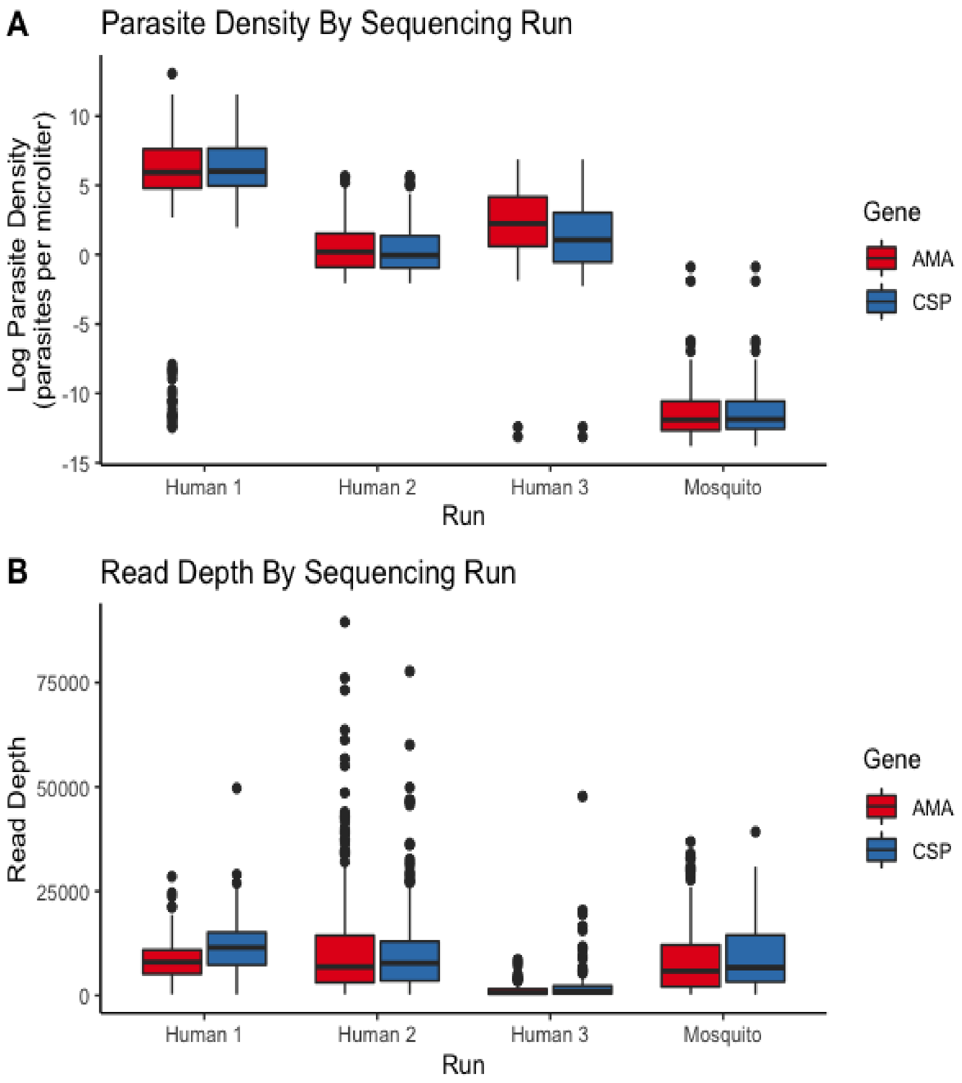


**Fig S 2:** Samples were split across four sequencing runs, three with human samples, and one with mosquito samples. (A) Parasite density of samples that were run on each of the four sequencing runs. Note that slightly different sets of samples were retained throughout filtering steps for *Pfcsp* and *Pfama1* genes, therefore both are shown. (B) Read depth at *Pfcsp* and *Pfama1* loci across samples that were run on each of the four sequencing runs.

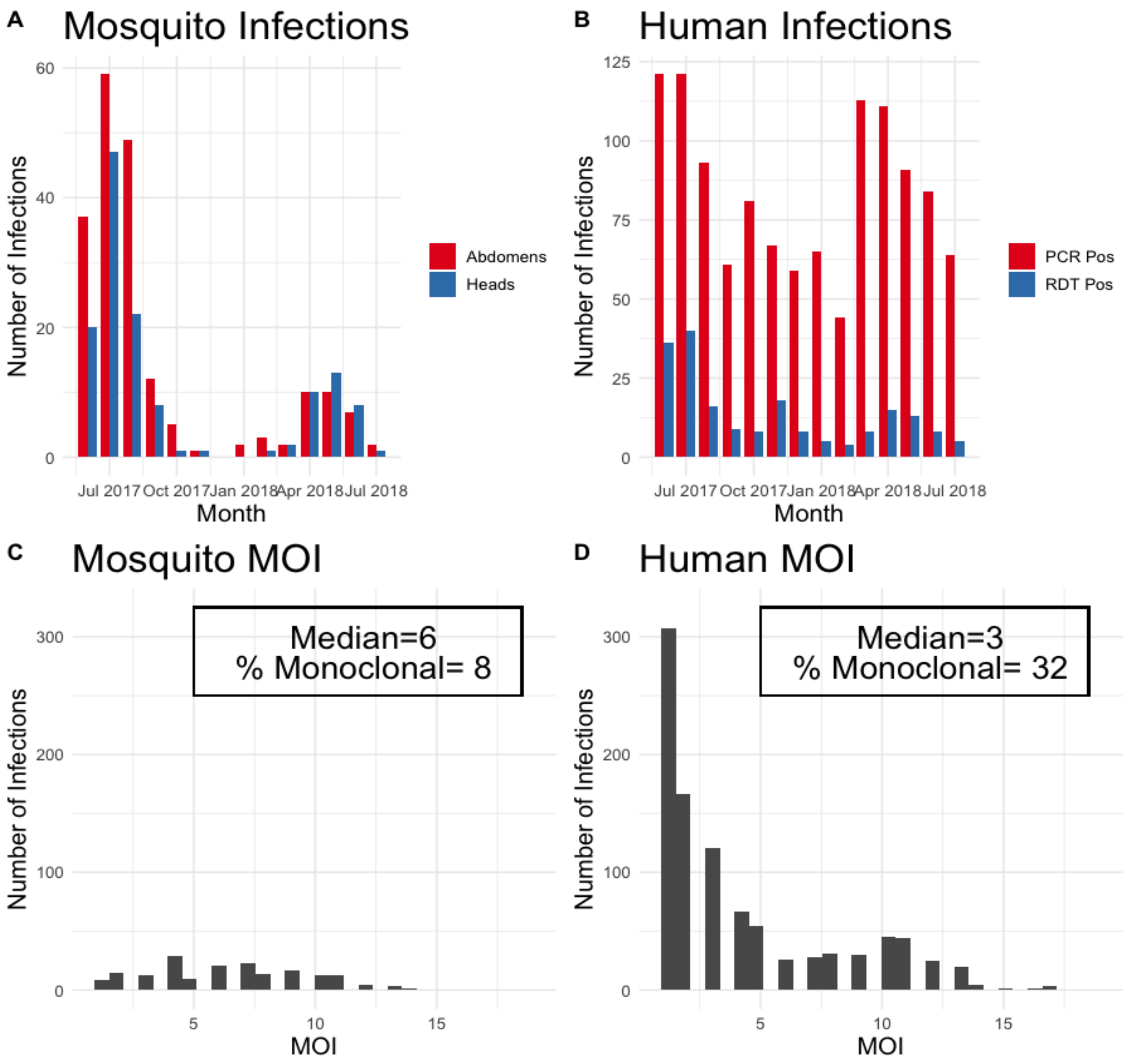


**Fig S 3:** Characteristics of infections within the mosquito and human populations over the study period, adapted from Sumner et al. [31]. A) Mosquito infections are separated by those detected in the abdomen or head and in B) humans by mode of detection via rapid diagnostic test (RDT) or polymerase chain reaction (PCR). The distribution of multiplicity of infection (MOI) values are shown in C) mosquito abdomens and D) humans as measured by the number of distinct Pfcsp haplotypes in each infection.

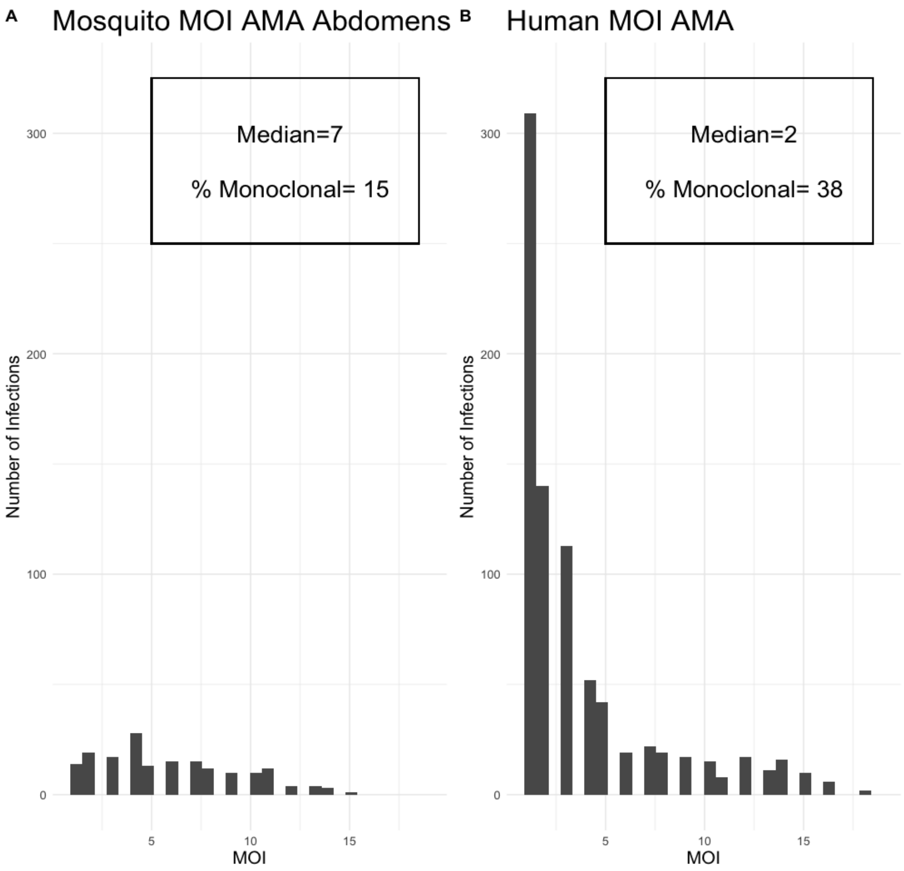


**Fig S 4:** Histograms of multiplicity of infection (MOI) in mosquito abdomens (A) and humans (B) as measured by the number of distinct *Pfama1* haplotypes in each infection.

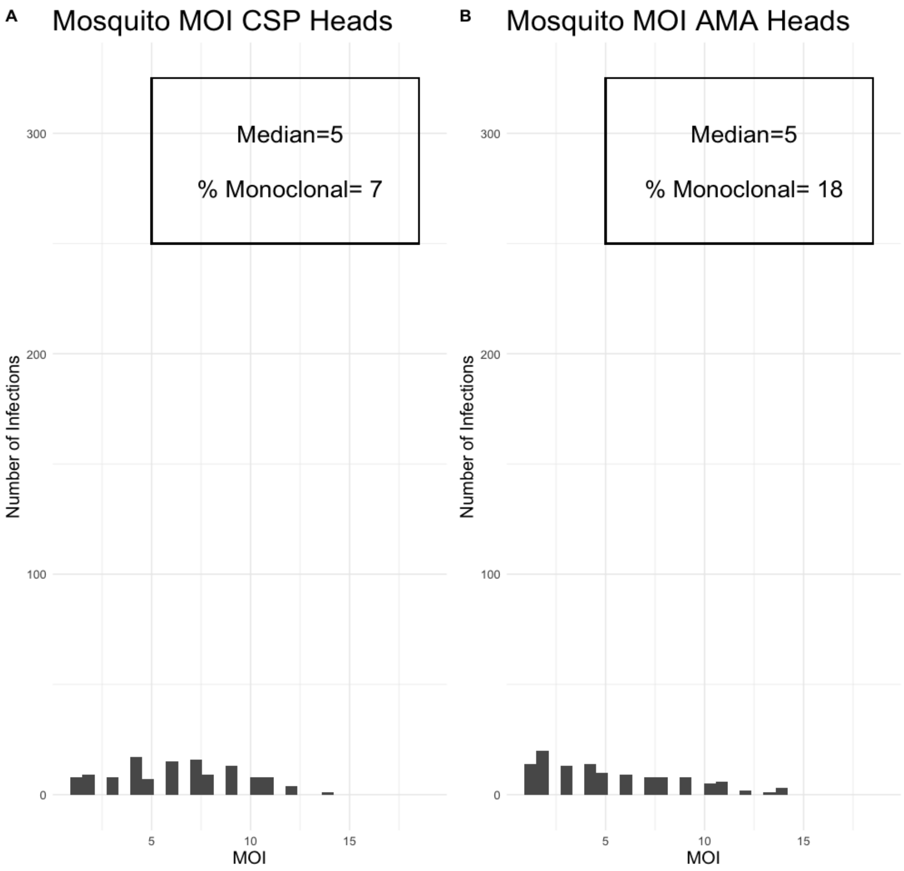


**Fig S 5:** Histograms of multiplicity of infection (MOI) in mosquito heads measured by the number of distinct *Pfcsp* haplotypes (A) and the number of distinct *Pfama1* (B) in each infection.

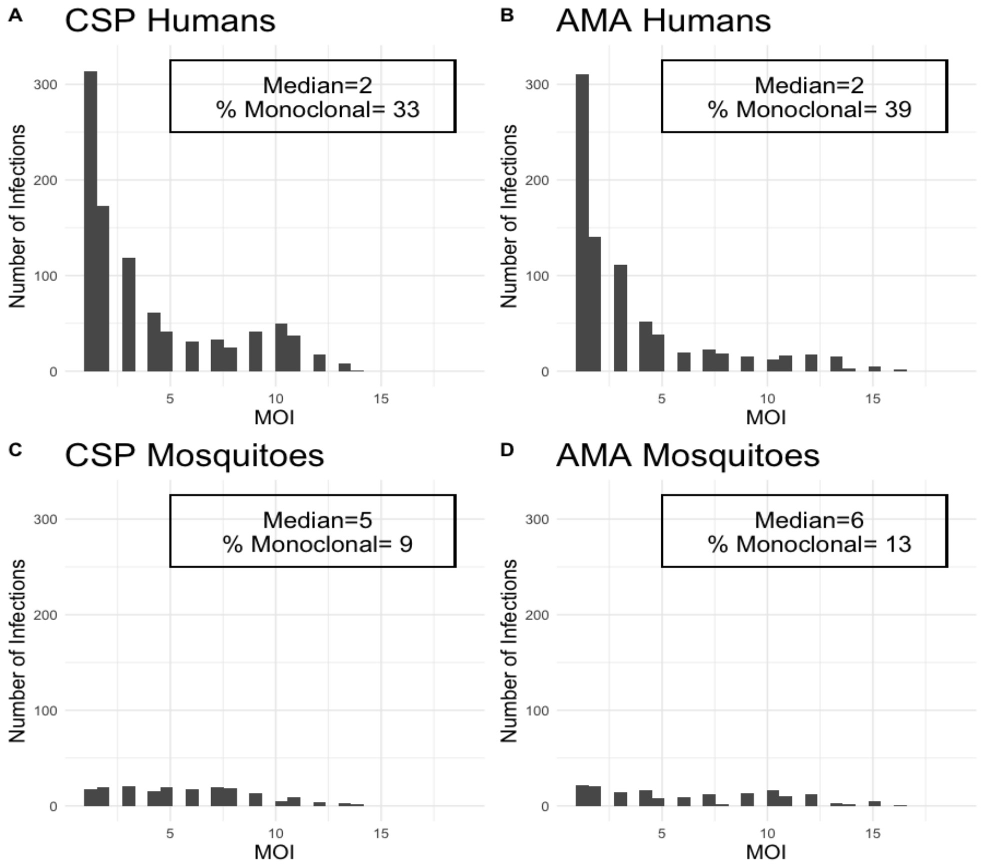


**Fig S 6:** Histograms of MOIs calculated by counting the number of unique haplotypes in infections, where common haplotypes (those with a population frequency above the population median frequency) only are considered. These are measured separately at the *Pfcsp* and *Pfama1* loci in humans (A, B) and mosquito abdomens (C, D).

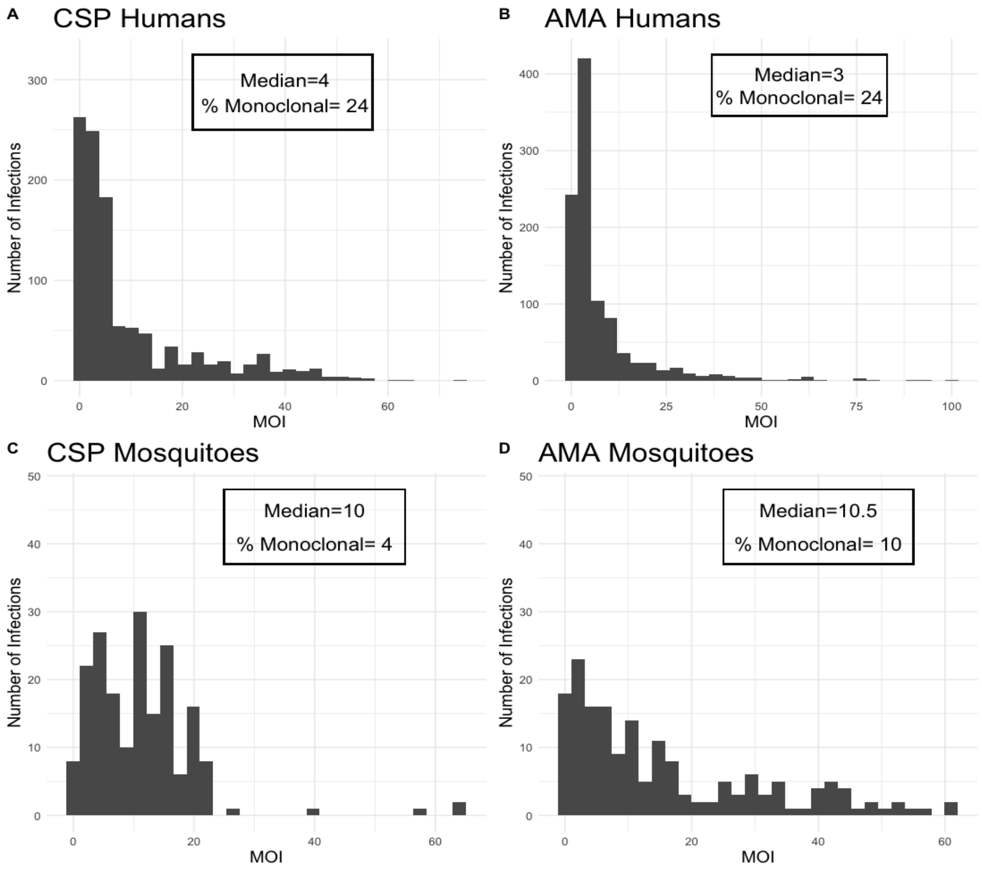


**Fig S 7:** Histograms of MOIs calculated by counting the number of unique haplotypes in infections, where all haplotypes from unfiltered sequencing data are considered. These are measured separately at the *Pfcsp* and *Pfama1* loci in humans (A, B) and mosquito abdomens (C, D).

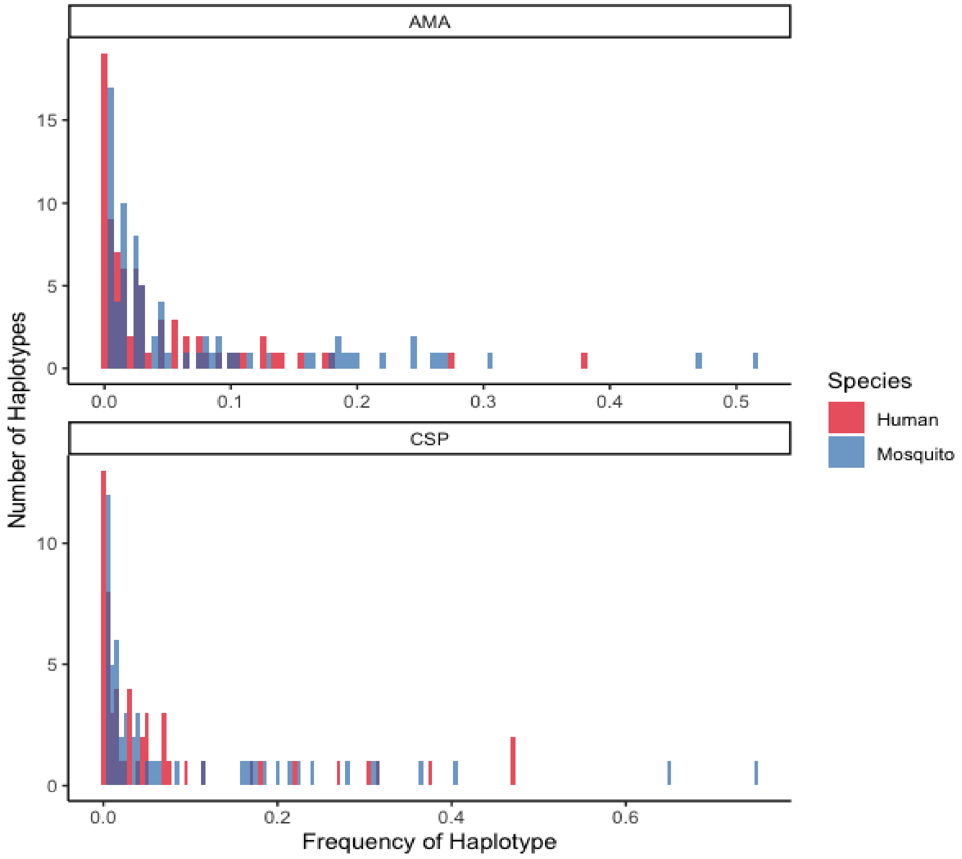


**Fig S 8:** Histograms of the frequency of haplotypes among human and mosquito infections for *Pfcsp* and *Pfama1* genes.

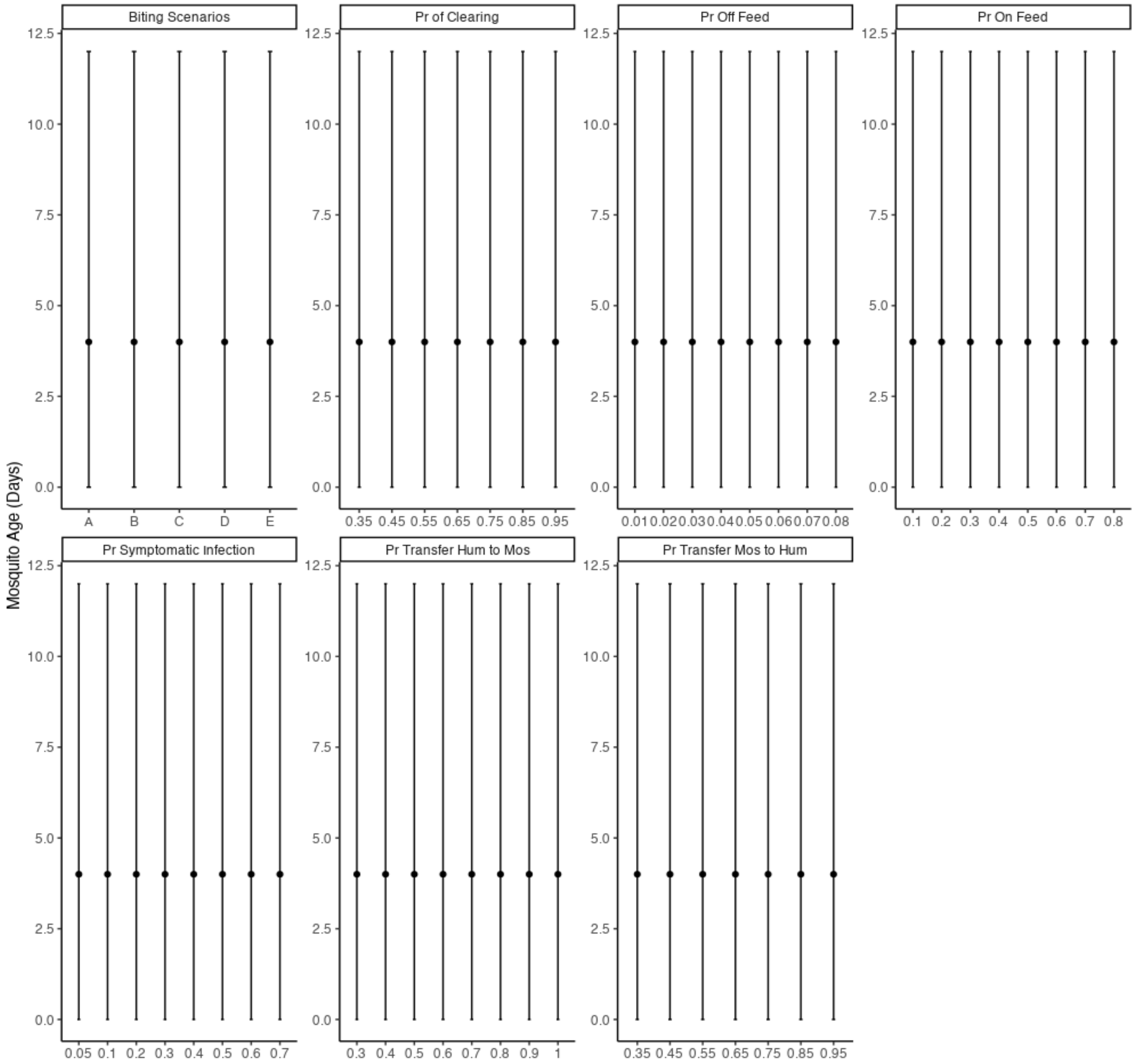


**Fig S 9:** Each square shows the median values and the $2.5$th and $97.5$th percentiles of mosquito age in days for 50 $365$ day long simulations on the Y axis. The X axis shows the values of various parameters under different simulation scenarios.

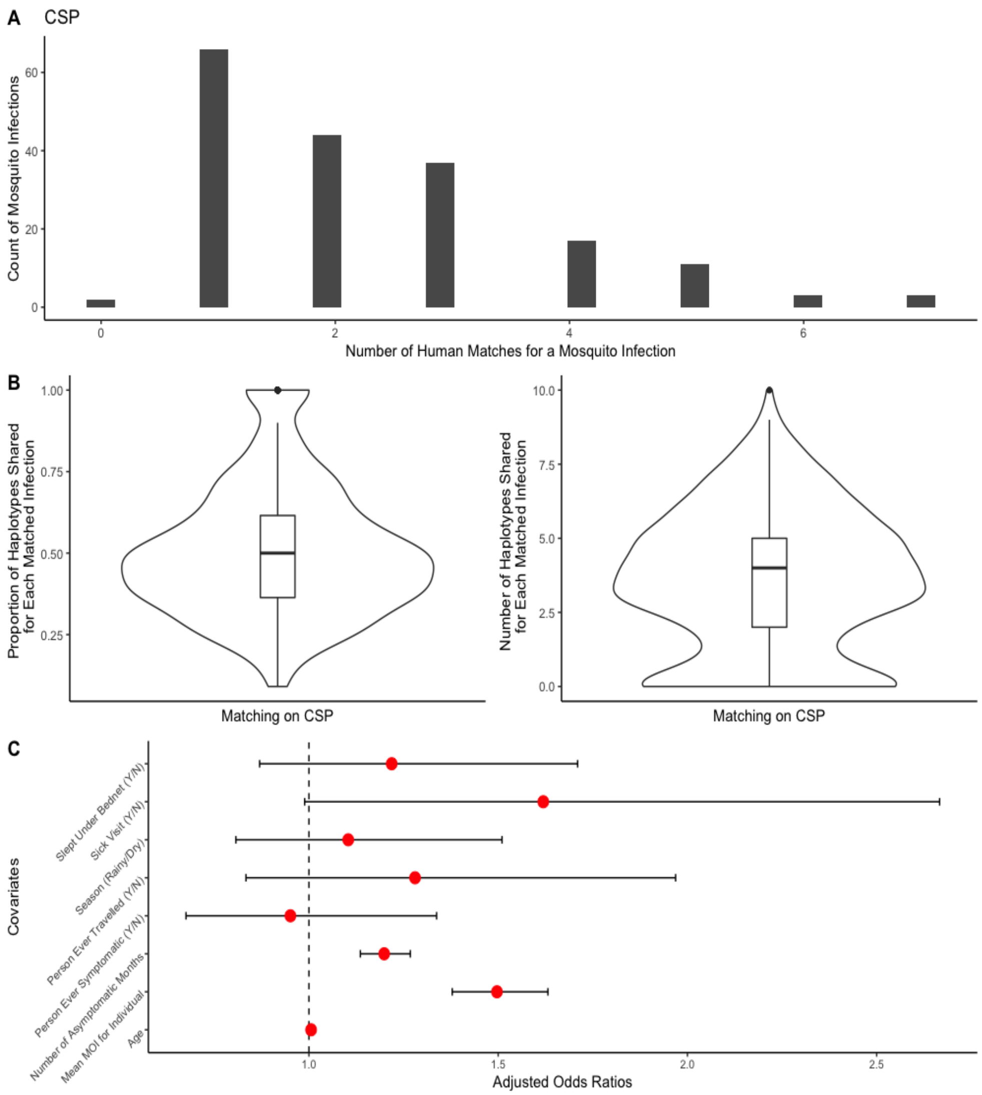


**Fig S 10:** A: Histograms of the number of human infections required to reconstitute mosquito infections obtained using the method that reconstitutes using *Pfcsp* haplotypes from mosquito abdomens. B: The proportion and number of haplotypes from each human infection also found in the matched mosquito's abdomen as a violin plot. C: Adjusted odds ratios with $95\%$ CIs from a multiple logistic regression with the probability of a human infection matching to at least one mosquito infection.

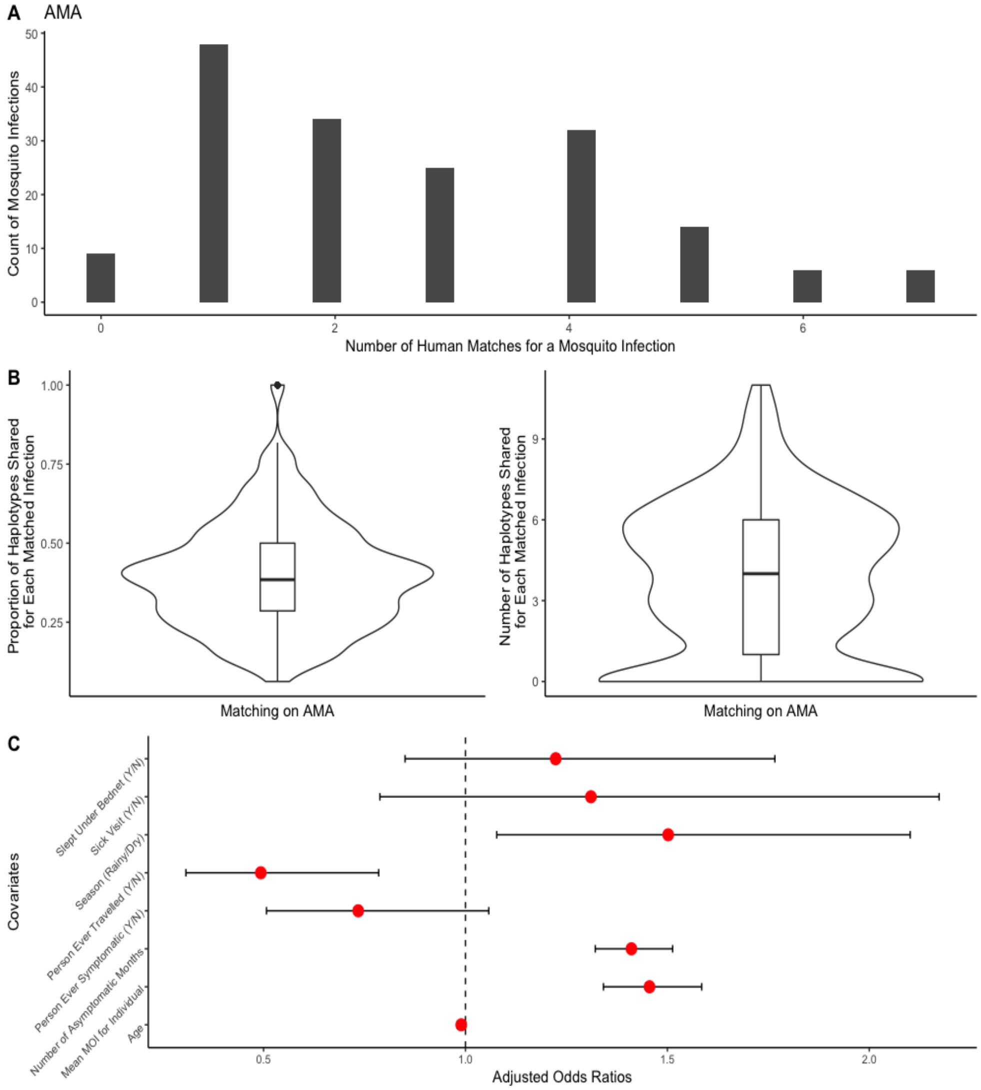


**Fig S 11:** A: Histograms of the number of human infections required to reconstitute mosquito infections obtained using the method that reconstitutes using *Pfama1* haplotypes from mosquito abdomens. B: The proportion and number of haplotypes from each human infection also found in the matched mosquito's abdomen as a violin plot. C: Adjusted odds ratios with $95\%$ CIs from a multiple logistic regression with the probability of a human infection matching to at least one mosquito infection.

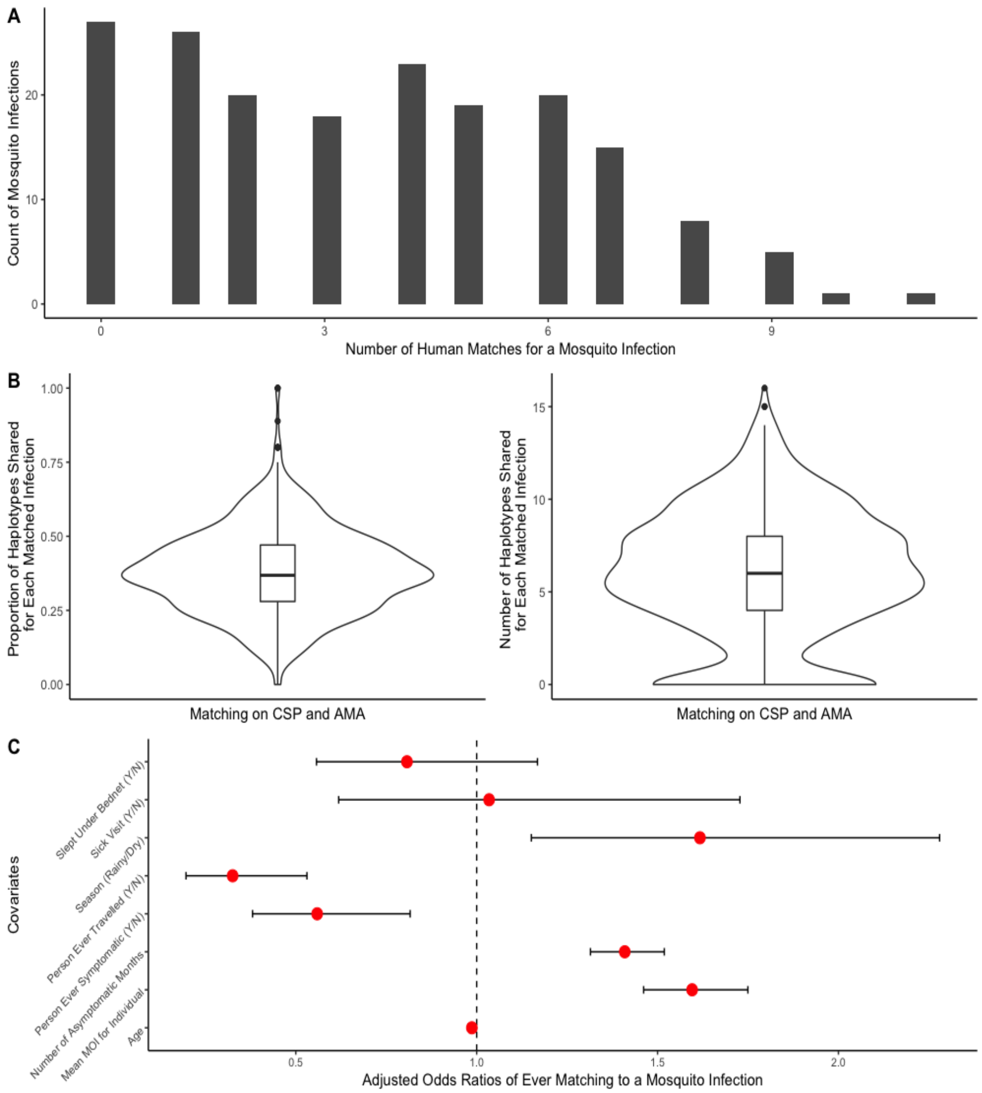


**Fig S 12:** (A) Histogram of the number of human infections required to reconstitute each mosquito infection obtained using the method that reconstitutes using common *Pfcsp* and *Pfama1* haplotypes from mosquito abdomens. Common *Pfcsp* and *Pfama1* haplotypes are those that have a frequency higher than the median frequency in both human and mosquito populations for that gene. (B) The proportion and number of haplotypes from each human infection also found in the matched mosquito's abdomen are on the y-axis. Matches are determined using the method that reconstitutes using both *Pfcsp* and *Pfama1* haplotypes from mosquito abdomens using only common haplotypes. (C) Adjusted odds ratios with $95\%$ CIs from a multiple logistic regression with the probability of a human infection matching to at least one mosquito infection using the method that reconstitutes mosquito infections using common *Pfcsp* and *Pfama1* haplotypes from mosquito abdomens.

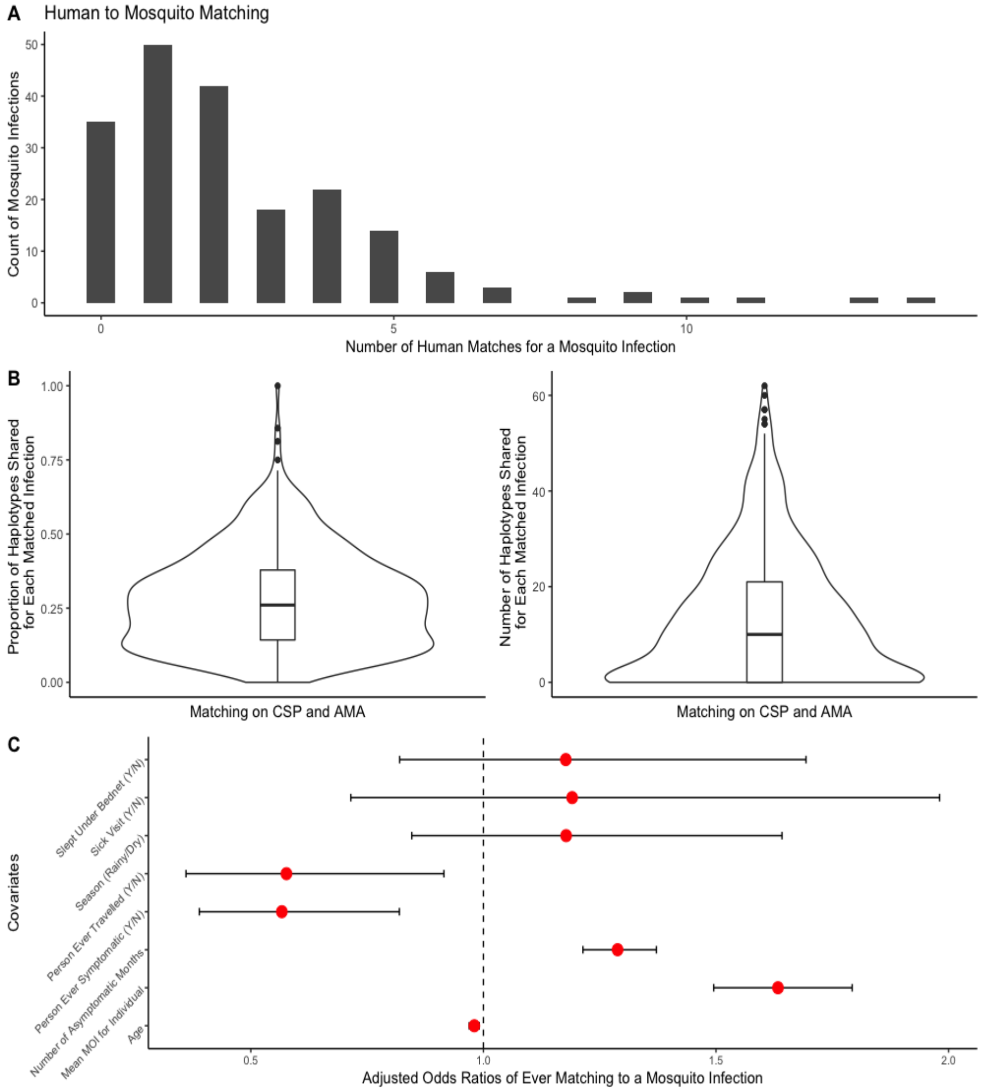


**Fig S 13:** (A) Histogram of the number of human infections required to reconstitute each mosquito infection obtained using the method that reconstitutes using unfiltered *Pfcsp* and *Pfama1* haplotypes from mosquito abdomens. (B) The proportion and number of haplotypes from each human infection also found in the matched mosquito's abdomen are on the y-axis. Matches are determined using the method that reconstitutes using both *Pfcsp* and *Pfama1* haplotypes from mosquito abdomens using unfiltered haplotypes. (C) Adjusted odds ratios with $95\%$ CIs from a multiple logistic regression with the probability of a human infection matching to at least one mosquito infection using the method that reconstitutes mosquito infections using unfiltered *Pfcsp* and *Pfama1* haplotypes from mosquito abdomens.

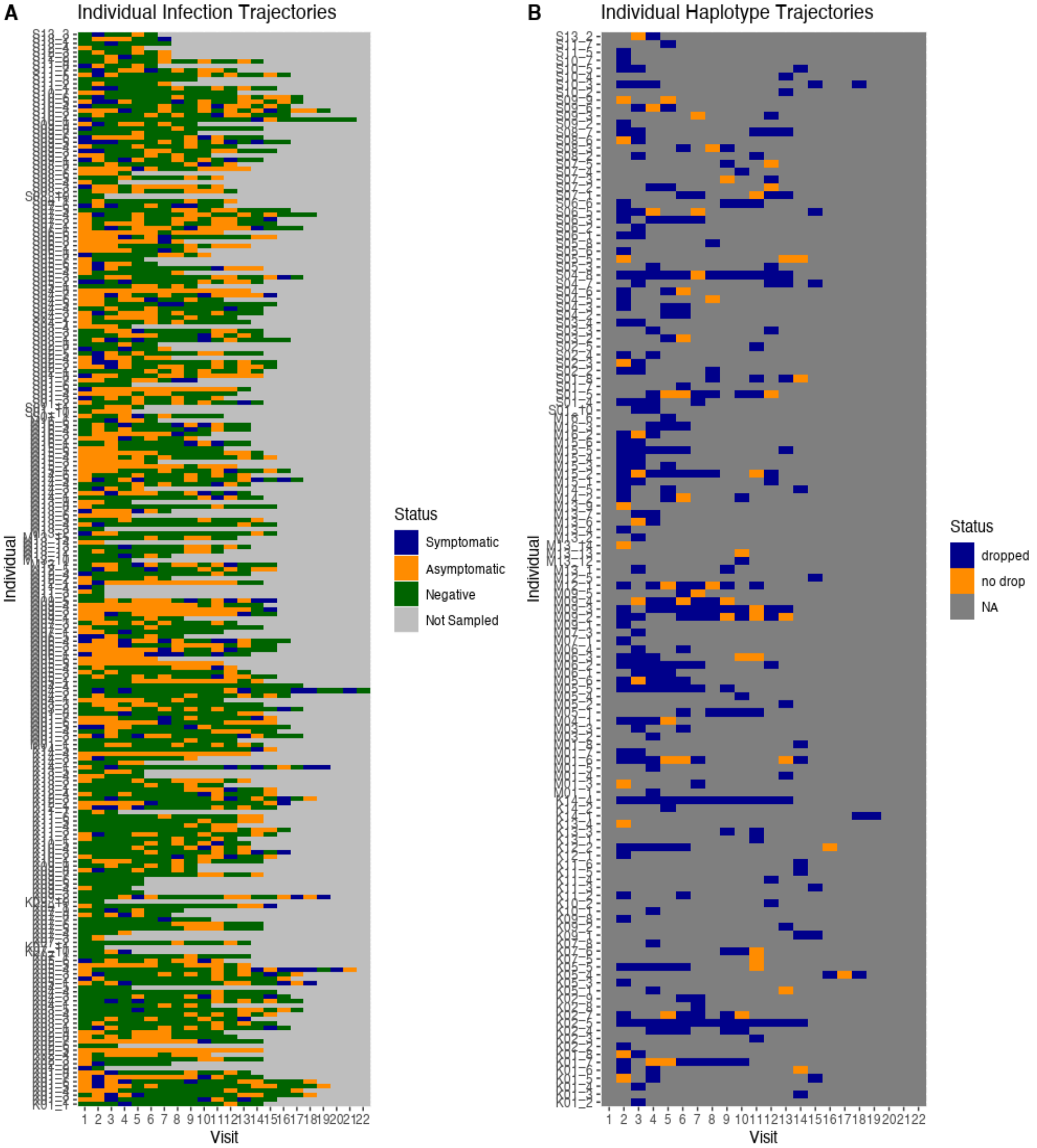


**Fig S 14:** (A) Individual infection trajectories showing whether an individual participant was symptomatic and positive by PCR, asymptomatic and positive by PCR or negative by PCR. Note that some participants had multiple missed visits throughout the study, and therefore were not sampled at all times, and that some participants had intermittent sick visits associated with symptomatic episodes. (B) Individual haplotype trajectories showing whether an individual has dropped haplotypes or not in consecutive sampled infections. NAs represent infections that were treated or times when the participant was not sampled.

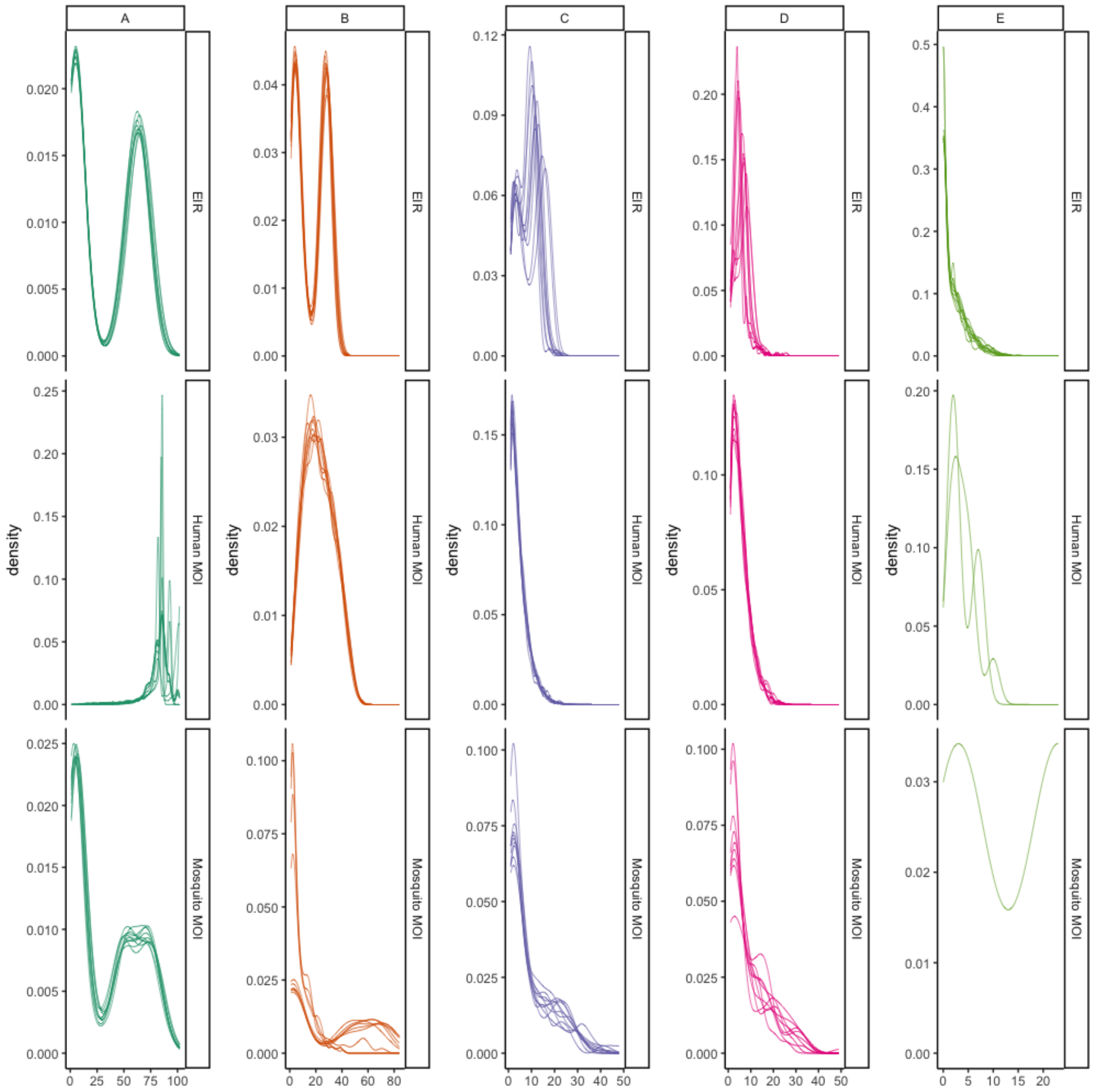


**Fig S 15:** Distribution of EIR, human and mosquito MOI for ten randomly selected simulations (of the 50 total simulations carried out for each scenario) under each biting scenario (A,B,C,D, and E). Note that under scenario E only two of the ten selected scenarios produced any humans with infections and none of the selected scenarios produced infected mosquitoes.

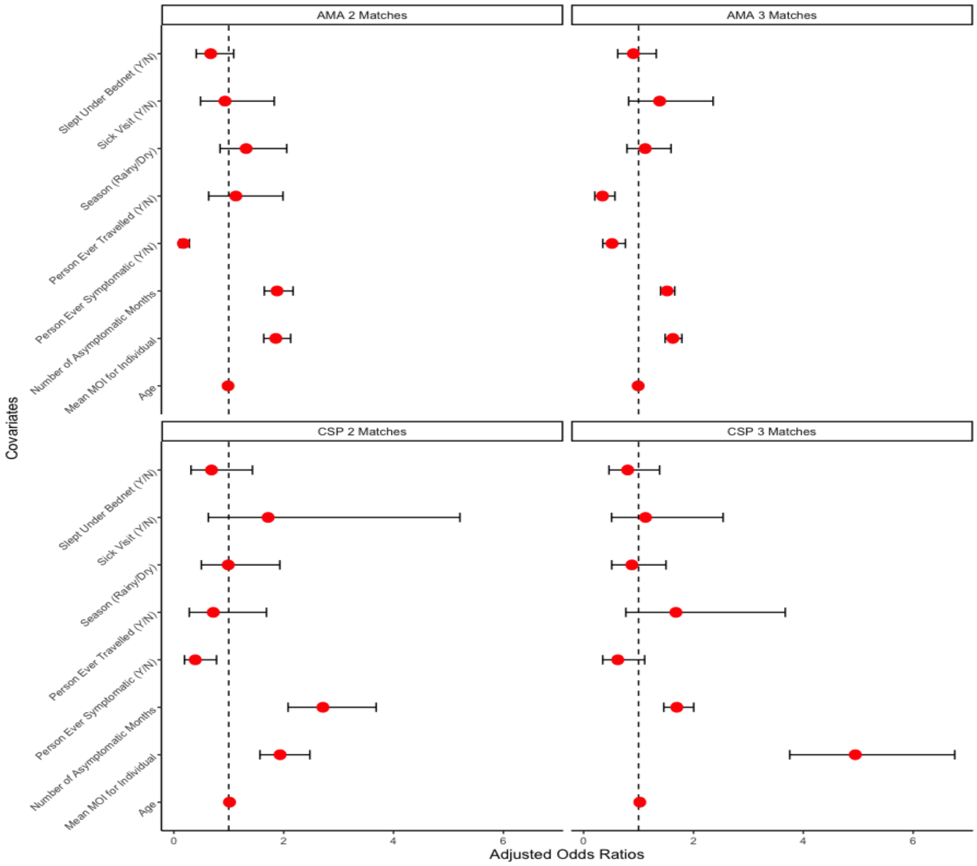


**Fig S 16:** Adjusted odds ratios with $95\%$ CIs from logistic regression models with the probability of a human infection matching to at least one mosquito infection using the method that reconstitutes mosquito infections by considering human infections that contain either 2 or 3 *Pfcsp* or *Pfama1* haplotypes found in the mosquito's abdomen.

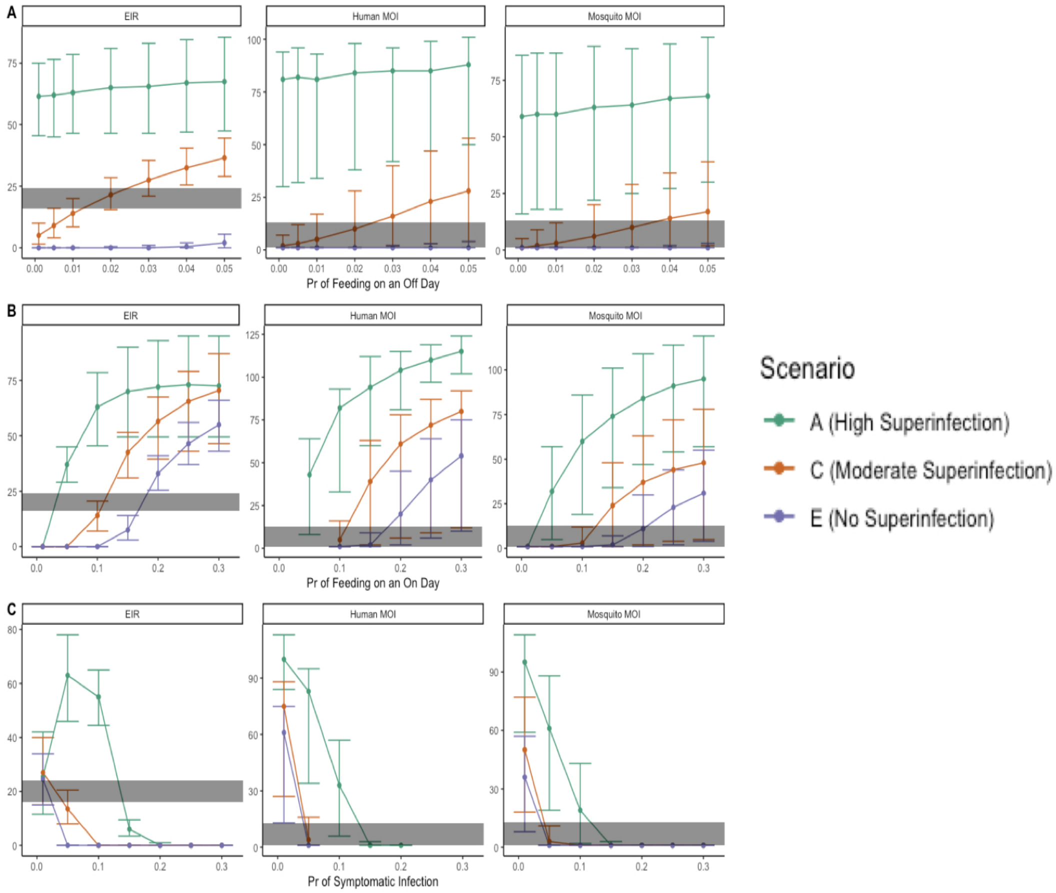


**Fig S 17:** The sensitivity of entomological inoculation rate (EIR), human and mosquito multiplicity of infection (MOI) under different simulation parameters and biting scenarios. Each square shows the median value and the 2.5th and 97.5th percentile of EIR, human MOI, or mosquito MOI across all simulations. We varied across parameters including (A) probability of feeding during an off day; (B) probability of feeding during an on day ; and (C) probability of a symptomatic infection. We show these results for three biting scenarios representing high probability of superinfection (Scenario A), medium probability of superinfection (Scenario C), and no probability of superinfection (Scenario E). The gray band represents the 2.5th and 97.5th percentile of the human and mosquito MOIs observed in the data and the range of published EIR estimates for the cohort area.

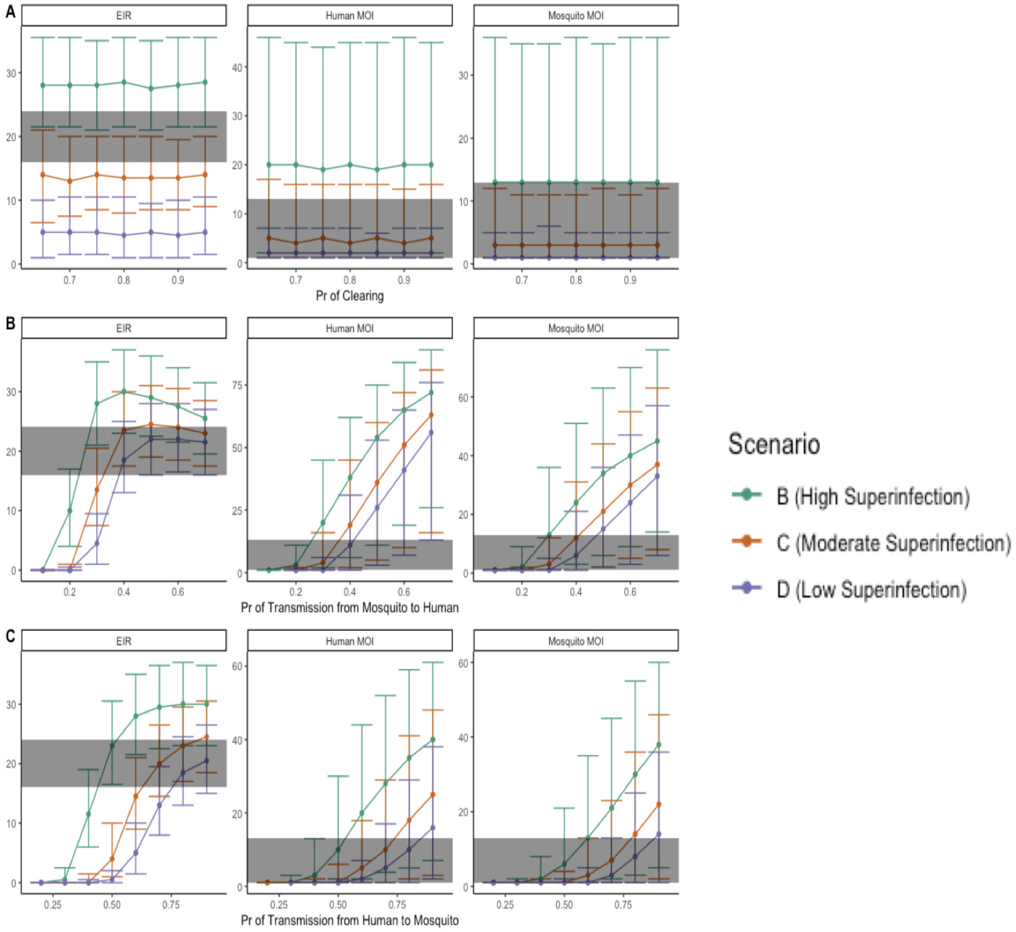


**Fig S 18:** The sensitivity of entomological inoculation rate (EIR), human and mosquito multiplicity of infection (MOI) under different simulation parameters and biting scenarios. Each square shows the median value and the 2.5th and 97.5th percentile of EIR, human MOI, or mosquito MOI across all simulations. We varied across parameters including (A) probability of haplotype clearance after 30 days; (B) probability of a haplotype being transferred from a mosquito to a human in an infectious bite;and (C) probability of a haplotype being transferred from a human to a mosquito in an infectious bite. We show these results for three biting scenarios representing high probability of superinfection (Scenario B), medium probability of superinfection (Scenario C), and low probability of superinfection (Scenario D). The gray band represents the 2.5th and 97.5th percentile of the human and mosquito MOIs observed in the data and the range of published EIR estimates for the cohort area.

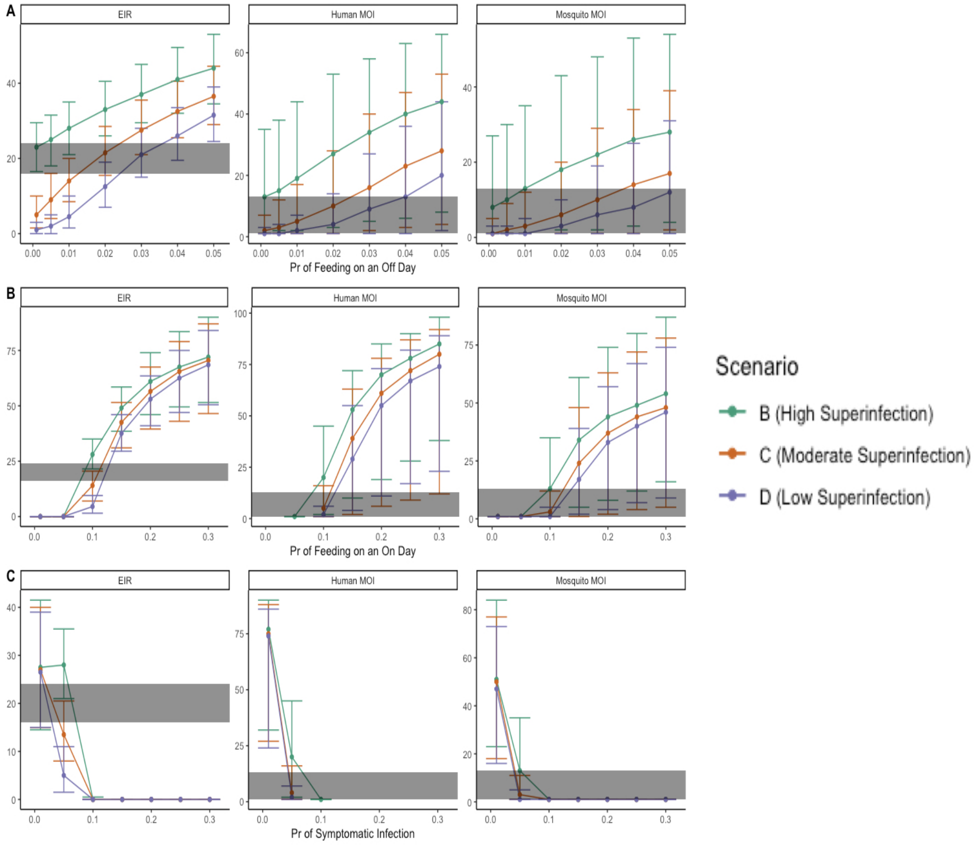


**Fig S 19:** The sensitivity of entomological inoculation rate (EIR), human and mosquito multiplicity of infection (MOI) under different simulation parameters and biting scenarios. Each square shows the median value and the 2.5th and 97.5th percentile of EIR, human MOI, or mosquito MOI across all simulations. We varied across parameters including (A) probability of feeding during an off day; (B) probability of feeding during an on day ; and (C) probability of a symptomatic infection. We show these results for three biting scenarios representing high probability of superinfection (Scenario B), medium probability of superinfection (Scenario C), and low probability of superinfection (Scenario D). The gray band represents the 2.5th and 97.5th percentile of the human and mosquito MOIs observed in the data and the range of published EIR estimates for the cohort area.

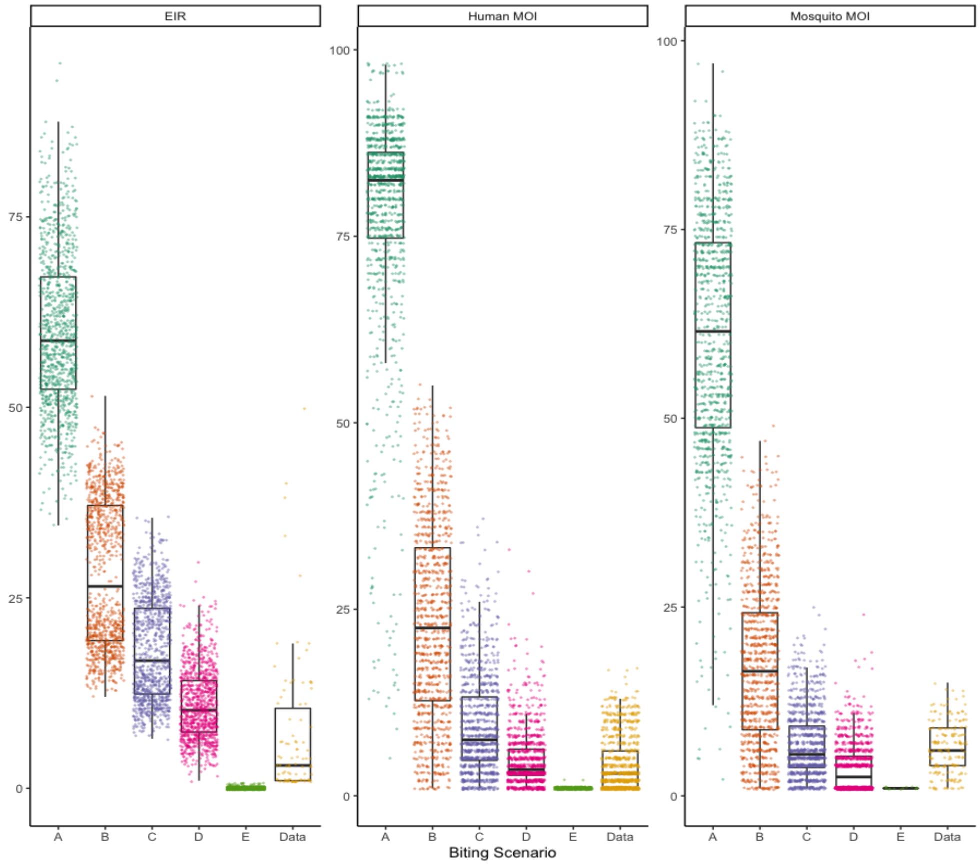


**Fig S 20:** Entomological inoculation rate (EIR), and MOI under different mosquito biting patterns under simulations where 20% of the population is designated more likely to be bitten. Boxplots of EIR, human MOI, or mosquito MOI across simulations by biting scenario using the baseline values of all other simulation parameters (probability of haplotype clearance after 30 days; probability of a haplotype being transferred from a mosquito to a human in an infectious bite; probability of a haplotype being transferred from a human to a mosquito in an infectious bite; probability of feeding during an off day; probability of feeding during an on day ; and probability of a symptomatic infection) for each simulation scenario. A random subset of 1000 points from the 50 simulation scenarios run for each biting pattern (A-E) are shown, all data points are shown. EIR was calculated from the data by counting the number of infectious bites each person received during the study period. Here, a person is deemed to have an infectious bite from a mosquito if they are identified as a match using the parsimonious matching method described in Section 2.3. Published estimates of EIR for this region range from 16 to 24 [40,41].

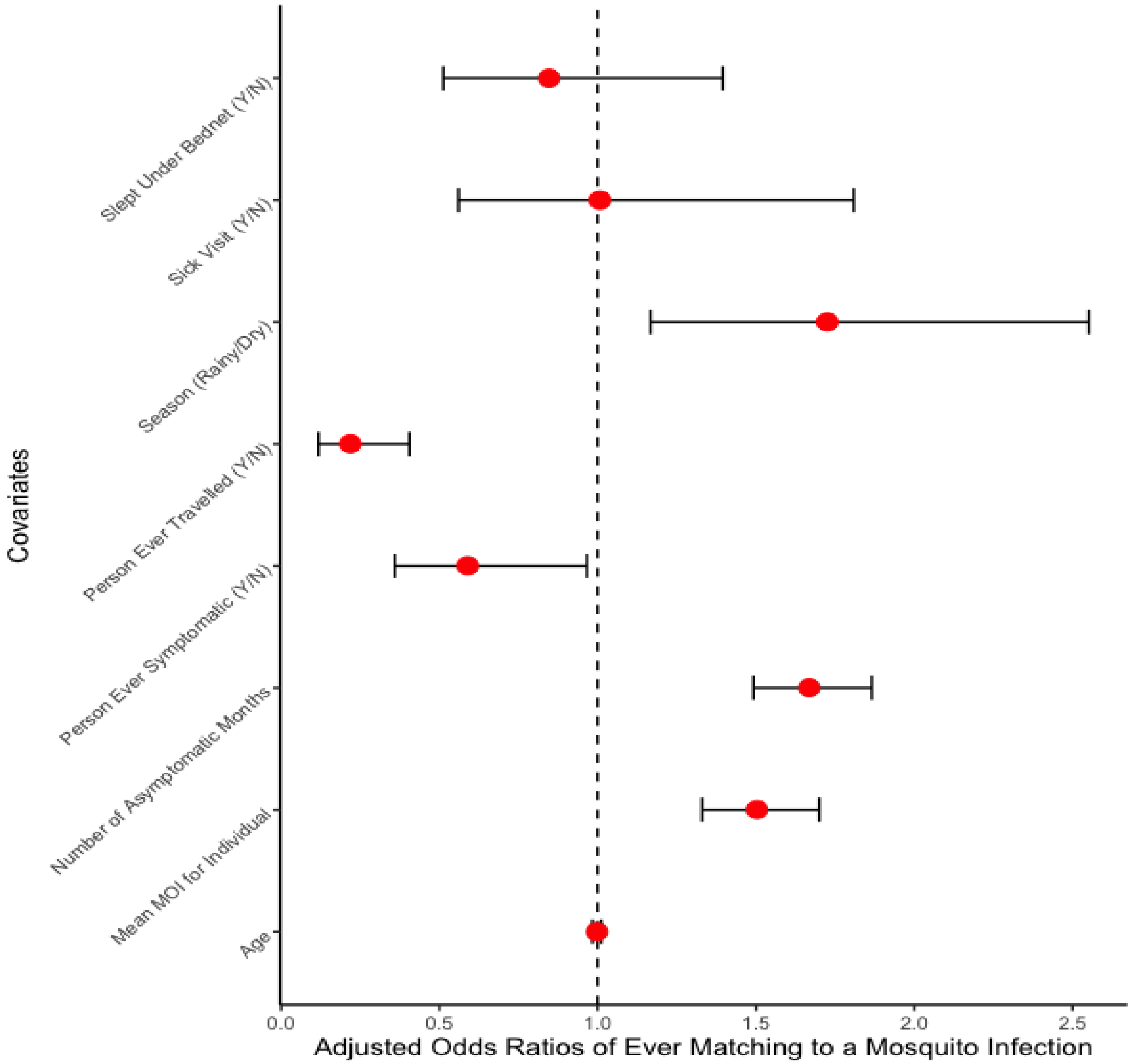


**Fig S 21:** Adjusted odds ratios with $95\%$ CIs from a multilevel logistic regression with the probability of a human infection matching to at least one mosquito infection using the method that reconstitutes mosquito infections using filtered *Pfcsp* and *Pfama1* haplotypes from mosquito abdomens. A random intercept for each household is used as a random effect. The conditional intra-class correlation coefficient (adjusting for the listed covariates) for this model is estimated to be $0.247$.

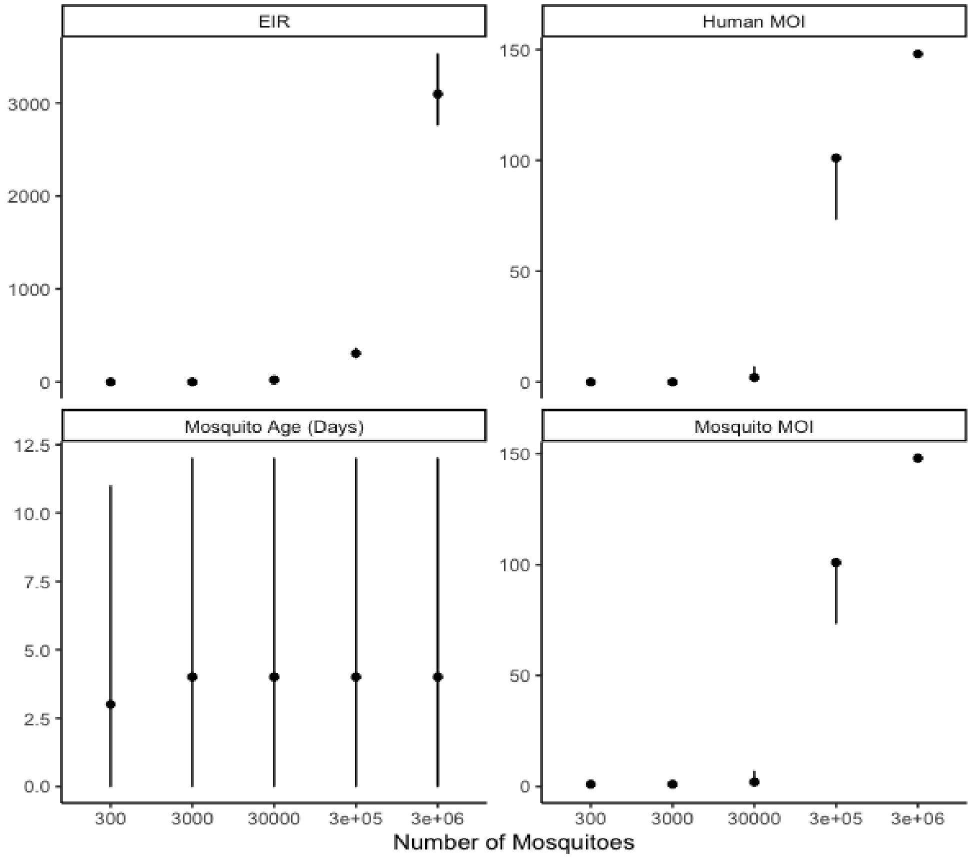


**Fig S 22:** Plot assessing the sensitivity of transmission parameters (entomological inoculation rate (EIR) and human and mosquito multiplicity of infection (MOI)) under different numbers of mosquitoes. Each square shows the median values and the $2.5$th and $97.5$th percentiles of EIR, mosquito age, human MOI, or mosquito MOI for one $365$ day long simulation on the Y axis. The X axis shows the number of mosquitoes.
